# Robust axis elongation by Nodal-dependent restriction of BMP signaling

**DOI:** 10.1101/2023.06.19.545591

**Authors:** Alexandra Schauer, Kornelija Pranjic-Ferscha, Robert Hauschild, Carl-Philipp Heisenberg

## Abstract

Embryogenesis is brought about by the coordinated activities of different signaling pathways controlling cell fate specification and morphogenesis. In vertebrate gastrulation, both Nodal and BMP signaling play key roles in germ layer specification and morphogenesis, yet their interplay to coordinate embryo patterning with morphogenesis is still insufficiently understood. Here, we took a reductionist approach using zebrafish embryonic explants to study the coordination of Nodal and BMP signaling for embryo patterning and morphogenesis. We show that Nodal signaling not only triggers explant elongation by inducing mesendodermal progenitors but also by suppressing BMP signaling activity at the site of mesendoderm induction. Ectopic BMP signaling in the mesendoderm blocks cell alignment and oriented mesendoderm intercalations, key processes to drive explant elongation. Translating these *ex vivo* observations to the intact zebrafish embryo showed that, similar to explants, Nodal signaling renders the dorsal domain less sensitive towards BMP signaling to allow effective cell intercalations and thus robust embryonic axis elongation. These findings suggest a dual function of Nodal signaling in embryonic axis elongation by both inducing mesendoderm and maintaining low levels of BMP signaling activity in the dorsal portion of the mesendoderm.

## Introduction

Embryonic development relies on cell fate specification and tissue shape changes mediated by highly conserved signaling pathways. These pathways often display dual functions in both instructing embryo patterning and determining the morphogenetic capacity of cells (Myers *et al*, 2002b; Heisenberg & Solnica-Krezel, 2008; Rogers & Schier, 2011; Briscoe & Small, 2015; Gilmour *et al*, 2017; Chan *et al*, 2017; Sagner & Briscoe, 2017; Pinheiro & Heisenberg, 2020; Lenne *et al*, 2021; Valet *et al*, 2022; Bailles *et al*, 2022). In line with this, cell fate specification is tightly linked with the emergence of distinct cellular behaviors, such as cell migration and intercalation, collectively leading to large-scale reorganization of tissues (Keller *et al*, 2003; Solnica-Krezel, 2005; Solnica-Krezel & Sepich, 2012; Chan *et al*, 2017; Gilmour *et al*, 2017; Hannezo & Heisenberg, 2019; Collinet & Lecuit, 2021; Valet *et al*, 2022). Thus, to robustly and reproducibly pattern and shape the embryo, cell fate specification and morphogenesis have to be closely coordinated in both space and time.

A hallmark of vertebrate gastrulation, the first major morphogenetic process in embryogenesis leading to the formation of the different germ layers (ectoderm, mesoderm and endoderm), is the elongation of the embryonic body along its anterior-posterior (AP) axis, involving both large-scale cell rearrangements and cell fate specification (Solnica-Krezel & Sepich, 2012). During zebrafish gastrulation, AP body axis elongation is driven by highly conserved convergence and extension (C&E) movements (Tada & Heisenberg, 2012; Williams & Solnica-Krezel, 2020a). These C&E movements are comprised of distinct cell behaviors, with cells undergoing both medially-oriented intercalations (mediolateral intercalations and planar medial intercalations (Yin *et al*, 2008; Glickman *et al*, 2003)) in dorsal positions of the gastrula and collective cell migration in anterior and ventrolateral domains, the combination of which is supposed to drive body axis elongation (Tada & Heisenberg, 2012; Williams & Solnica-Krezel, 2020a). Concomitant with AP body axis elongation, the tissues undergoing C&E movements become patterned along their AP and dorsal-ventral (DV) axes by the graded activity of the conserved transforming growth factor β (TGFβ) signals Nodal and Bone Morphogenetic Protein (BMP), which both have been implicated in regulating axis patterning as well as morphogenesis during zebrafish gastrulation (Myers *et al*, 2002b; Schier & Talbot, 2005; Heisenberg & Solnica-Krezel, 2008; Zinski *et al*, 2018; Pinheiro & Heisenberg, 2020; Williams & Solnica-Krezel, 2020a; Hill, 2022; Pinheiro *et al*, 2022). While the processes setting up the graded activity domains of these signals, also termed ‘morphogens’, and their capacity to control cell fate specification and cell behavior have been studied for some time (Myers *et al*, 2002b; Schier & Talbot, 2005; Heisenberg & Solnica-Krezel, 2008; Zinski *et al*, 2018; Rogers & Müller, 2019; Pinheiro & Heisenberg, 2020; Williams & Solnica-Krezel, 2020a; Hill, 2022; Pinheiro *et al*, 2022), much less is still known about how their combined activities are coordinated in space and time.

Seminal work in *Xenopus* has demonstrated that cultured gastrula explants containing the dorsal blastopore lip or prospective animal ectoderm treated with the TGFβ signal Activin can undergo medio-lateral cell intercalations leading to axis elongation (Keller *et al*, 1985; Symes & Smith, 1987; Keller & Danilchik, 1988; Thomsen *et al*, 1990; Sokol & Melton, 1991; Wilson & Keller, 1991; Shih & Keller, 1992a, 1992b; Keller *et al*, 1992; Green *et al*, 2004; Ninomiya *et al*, 2004). Interestingly, *Xenopus* explant elongation depends on proper AP patterning of the mesendoderm (Ninomiya *et al*, 2004), and the specific combination of different factors inducing mesendoderm specification (Green *et al*, 1990; Cunliffe & Smith, 1992; Howard & Smith, 1993; Graff *et al*, 1994), suggesting a close link between morphogen signaling, tissue patterning and axis elongation also *in vitro*. More recently, mammalian embryonic stem cell-derived 3D *in vitro* models of gastrulation (gastruloids) have been shown to elongate, suggesting that axis elongation is a conserved feature of these systems (van den Brink *et al*, 2014; Turner *et al*, 2017; Beccari *et al*, 2018; Moris *et al*, 2020; Anlaş *et al*, 2021; Xu *et al*, 2021). Similar to *Xenopus* explants, the signaling regime during gastruloid formation affects their elongation capacity and shape (van den Brink *et al*, 2014; Turner *et al*, 2017; Anlaş *et al*, 2021; Underhill & Toettcher, 2023; Hennessy *et al*, 2023). This suggests that also in these reduced stem cell assay systems of gastrulation, the appropriate spatiotemporal organization of morphogen signaling is a key factor to control axis elongation.

Here, we have used zebrafish embryonic explants to ask how the spatial organization of morphogen signaling domains affects axis elongation. We found that explant elongation is predominantly driven by oriented mesendodermal cell intercalations, which can only occur when BMP signaling levels are sufficiently low within the mesendoderm. Nodal signaling, in addition to its role in triggering mesendoderm induction, maintains such low levels of BMP signaling at the site of explant elongation by suppressing BMP signaling, a critical function also required in the intact embryo for robust body axis elongation.

## Results

### Explant elongation during gastrulation is driven by mesendoderm cell intercalation behavior

Body axis elongation in zebrafish embryos during gastrulation is the result of the concerted action of various distinct cell behaviors characteristic for different germ layer cell identities that are set up by morphogen gradients within the embryo (Myers *et al*, 2002b; Heisenberg & Solnica-Krezel, 2008; Tada & Heisenberg, 2012; Zinski *et al*, 2018; Williams & Solnica-Krezel, 2020a). Zebrafish embryonic explants, similar to other *in vitro* models of gastrulation (Anlas & Trivedi, 2021; Steventon *et al*, 2021; Emig & Williams, 2022), undergo elongation reminiscent of embryonic axis extension in the intact embryo upon local Nodal signaling activation (Fig. 1A,B, Fig. S1A) (Xu *et al*, 2014; Fulton *et al*, 2020; Williams & Solnica-Krezel, 2020b; Schauer *et al*, 2020; Cheng *et al*, 2023). However, the underlying cell rearrangements and contribution of different cell types in this process are not yet fully understood. To address this, we first analyzed which progenitor types drive the elongation movement. In the initial phase of explant elongation, the extended part of the explant was predominantly composed of mesendodermal tissues, as identified by the expression of the pan-mesendodermal marker *sebox* (Poulain & Lepage, 2002; Ruprecht *et al*, 2015) (Fig. 1C,D, S1B,C). Moreover, during explant elongation the mesendodermal tissue within the explant extension was changing its shape drastically, elongating in the direction of the extension and narrowing perpendicular to it (Fig. 1E,F, Fig. S1D, Supplementary Video 1), while keeping its total area largely constant (Fig. S1E). This is consistent with suggestions that whole explant elongation is driven by convergence and extension movements (C&E movements) rather than oriented growth (Xu *et al*, 2014; Williams & Solnica-Krezel, 2020b; Fulton *et al*, 2020; Schauer *et al*, 2020; Cheng *et al*, 2023). Lowering Nodal signaling activity and concomitantly the amount of mesendoderm formed in the explants reduces the extent of explant elongation (Fig. S1F,F’), further supporting a link between mesendoderm C&E movements and overall explant elongation.

**Figure 1.**
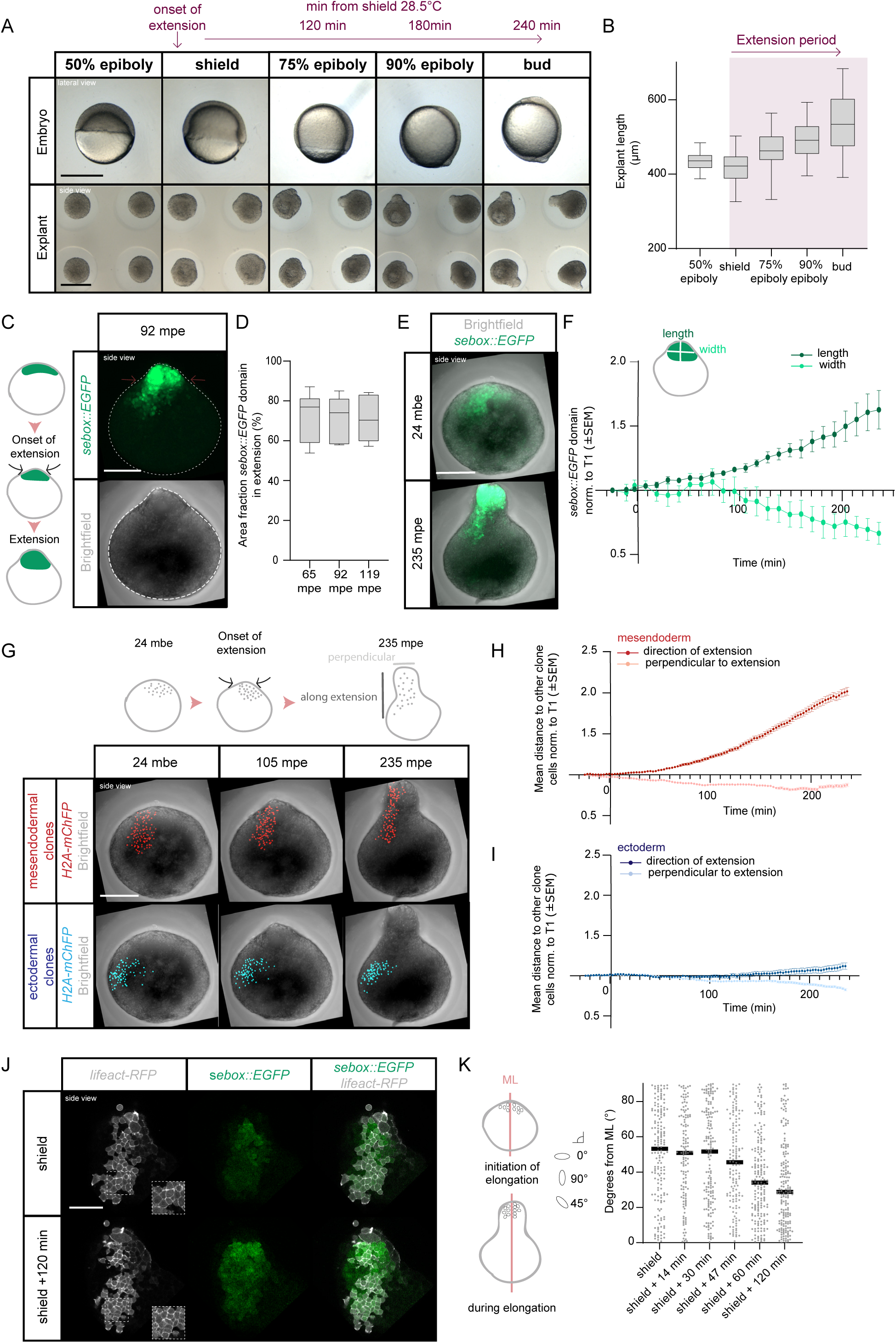
Mesendoderm morphogenesis and cell dispersal during blastoderm explant elongation. (**A**) Single-plane brightfield images of embryos (lateral view) and blastoderm explants (side view) at consecutive stages from 50% epiboly to bud stage. The timeline on top of the images indicates the onset of blastoderm explant extension and the time between the indicated developmental stages at 28.5°C. (**B**) Blastoderm explant length at consecutive stages from 50% epiboly to bud stage (n=45, N=3). (**C**) Maximum intensity projection of fluorescence (top) and brightfield (bottom) images (side views) of blastoderm explants obtained from Tg(*sebox::EGFP*) embryos expressing EGFP (green) in mesendoderm progenitors at 92 minutes post explant elongation onset (mpe). (**D**) Area fraction of EGFP expression (mesendoderm progenitors) within the extension of explants obtained from Tg(*sebox::EGFP*) embryos at 65 (n=7, N=7), 92 (n=7, N=7) and 119 mpe (n=7, N=7). (**E**) Maximum intensity projection of fluorescence/brightfield images (side views) of blastoderm explants from Tg(*sebox::EGFP*) embryos expressing EGFP (green) in mesendoderm progenitors 24 min before the onset of explant elongation (mbe, top) and at 235 mpe (bottom). (**F**) Length and width of the EGFP expression domain in blastoderm explants from Tg(*sebox::EGFP*) embryos marking mesendoderm progenitors during explant elongation (n=6, N=6). Length is calculated along the direction of explant elongation and width is oriented perpendicular to the length. (**G**) Maximum intensity projection of fluorescence/brightfield images (side views) of blastoderm explants before explant elongation (24 mbe, left column), during explant elongation (105 mpe, middle column) and at the end of elongation (235 mpe, right column) showing clonally labeled cell nuclei in the mesendoderm (red) or ectoderm (blue). (**H,I**) Mesendodermal (H) and ectodermal (I) clone dispersal parallel and perpendicular to the axis of explant elongation assessed by the mean distance of each cell in the clone to other clone cells during explant elongation (mesendoderm > 320 cells analyzed per time point, n=5, N=5; ectoderm > 240 cells analyzed per time point, n=5, N=5). (**J**) Single-plane high-resolution images of blastoderm explants (side views) obtained from Tg(*sebox::EGFP*) embryos expressing EGFP in mesendoderm progenitors at the onset of explant elongation (corresponding to embryonic shield stage) and during explant elongation (shield + 120 min). Mesendodermal cell clones within the explant extension are marked by *lifeact-RFP* (grey) expression. (**K**) Cell alignment assessed by the deviation (degrees) of the main cell extension axis from the main mediolateral (ML) explant axis at the onset of explant elongation (shield: 168 cells, n=5, N=5) and during explant elongation (shield + 14 min: 132 cells, n=5, N=5, shield + 30 min: 162 cells, n=5, N=5, shield + 47 min: 144 cells, n=5, N=5, shield + 60 min: 184 cells, n=5, N=5, shield + 120 min: 188 cells, n=5, N=5). Scale bars: 500 µm (A), 200 µm (C,E,G), 100 µm (J).

To gain insight into the cell rearrangements occurring within the mesendoderm during explant elongation, we analyzed the dispersal of *sebox::EGFP* positive-cells over time by generating cell clones of varying sizes and position within the mesendoderm expressing *H2A-mChFP* marking their nuclei (Fig. 1G,H). During explant elongation, the mean distance between clonal cells increased along the direction of the explant extension and decreased perpendicular to it (2.02-fold change in distance in direction of and 0.88-fold change in distance perpendicular to extension over 260 min) (Fig. 1G,H, Fig. S1G,G’, Supplementary Video 2). The position of mesendodermal progenitors relative to each other along the explant elongation axis remained largely unchanged during the elongation process as evidenced by their relative distance to the tip of the explant prior to the onset of extension being correlated with their final position (R^2^=0.6748, Fig. S1H). To determine whether this clonal behavior was the result of medially-oriented mesendodermal cell intercalations, similar to the situation in dorsal tissues of the intact embryo (Glickman *et al*, 2003; Yin *et al*, 2008), we analyzed the orientation of the longest cell axis relative to the explant mediolateral axis during explant elongation (Fig. 1J,K) (Williams & Solnica-Krezel, 2020b). At the onset of explant elongation (corresponding to embryonic shield stage), the cells appeared randomly oriented (Fig. 1J,K, Fig. S1I, Supplementary Video 3). However, during the subsequent elongation process, the longest cell axis became increasingly aligned with the mediolateral explant axis (Fig. 1J,K, Fig. S1I), consistent with the notion that these cells were undergoing coordinated cell alignment, a hallmark for medially-oriented cell intercalations (Glickman *et al*, 2003).

Interestingly extending our cell dispersal analysis to the ectoderm showed that ectodermal progenitors initially exhibited only very limited cell rearrangements (0.99-fold change in distance in direction of extension over 122 min compared to 1.2-fold change for mesendodermal progenitors) and that clone elongation in the direction of the extension was largely restricted to late stages of the elongation process (1.12-fold change in distance in direction of and 0.83-fold change in distance perpendicular over 260 min) (Fig. 1G,I, Fig. S1G,G’, Supplementary Video 4). Collectively, these findings suggest that explant elongation is predominantly driven by medially-oriented mesendoderm cell intercalations.

### BMP-mediated dorsoventral patterning determines the elongation capacity of explants

Given that mesendodermal cell rearrangements are important for explant elongation and that different mesendodermal cell fates display distinct morphogenetic capacities in the zebrafish embryo (Tada & Heisenberg, 2012; Williams & Solnica-Krezel, 2020a), we assessed the role of mesendodermal patterning for explant elongation. Taking advantage of the inherent variability in the induction of various mesodermal cell fates in explants (Schauer *et al*, 2020; Fulton *et al*, 2020), we first analyzed the relationship between explant elongation/length and the size of the expression domain of dorsal (*hgg* (Thisse *et al*, 1994), *flh* (Talbot *et al*, 1995)), paraxial (*papc*) (Yamamoto *et al*, 1998), ventrolateral (*tbx16*) (Griffin *et al*, 1998) and ventral (*drl*) (Herbomel *et al*, 1999) mesodermal markers at the end of explant elongation, corresponding to bud stage in intact embryos (Fig. S2A-C). While there was no clear correlation between the size of the expression domain of the tested markers and explant length (Fig. S2A), we noted that markers for the most ventral mesodermal cell fates tested were rarely expressed in explants (Fig. S2B,C). To test whether the induction of these most ventral mesodermal cell fates might interfere with explant elongation, we analyzed the effect of changes in BMP signaling levels, a key determinant for zebrafish dorsoventral (DV) patterning (Zinski *et al*, 2018), on explant elongation. To reduce or increase BMP signaling, we prepared explants from embryos treated with various positive and negative regulators of BMP signaling. We found that a reduction in BMP signaling levels by morpholino-mediated *bmp2b* knockdown (Lele *et al*, 2001a) or overexpression of a GFP-tagged version of the BMP antagonist Chordin (*chrd-GFP* (Hammerschmidt *et al*, 1996; Pomreinke *et al*, 2017)) in explants neither affected the frequency of explant elongation nor the average length of the extension (Fig. 2A-D, Fig. S2D-G). In contrast, increasing BMP signaling and thereby increasing the most ventral mesodermal cell fates by overexpression of *bmp2b* or *caAlk8* (a constitutively active BMP receptor (Bauer *et al*, 2001); Fig. S2B,C) severely reduced both the frequency of explant elongation (Fig. 2A-C, Fig. S2D-F) and the average length of the extension (Fig. 2A,D, Fig. S2D,G), accompanied by a reduced length/width ratio of the mesendoderm (Fig. 2E,F, Fig. S3A,B). This reduction in explant elongation frequency was dependent on the extent of BMP overactivation, as assessed by overexpressing different amounts of *bmp2b* and *caAlk8* (Fig. 2B, S2E), and required the activity of the transcriptional regulator *Smad5* (Fig. S3C-F) (von der Hardt *et al*, 2007). This suggests that overactivation, but not abrogation of BMP signaling affects mesendoderm-driven explant elongation.

**Figure 2.**
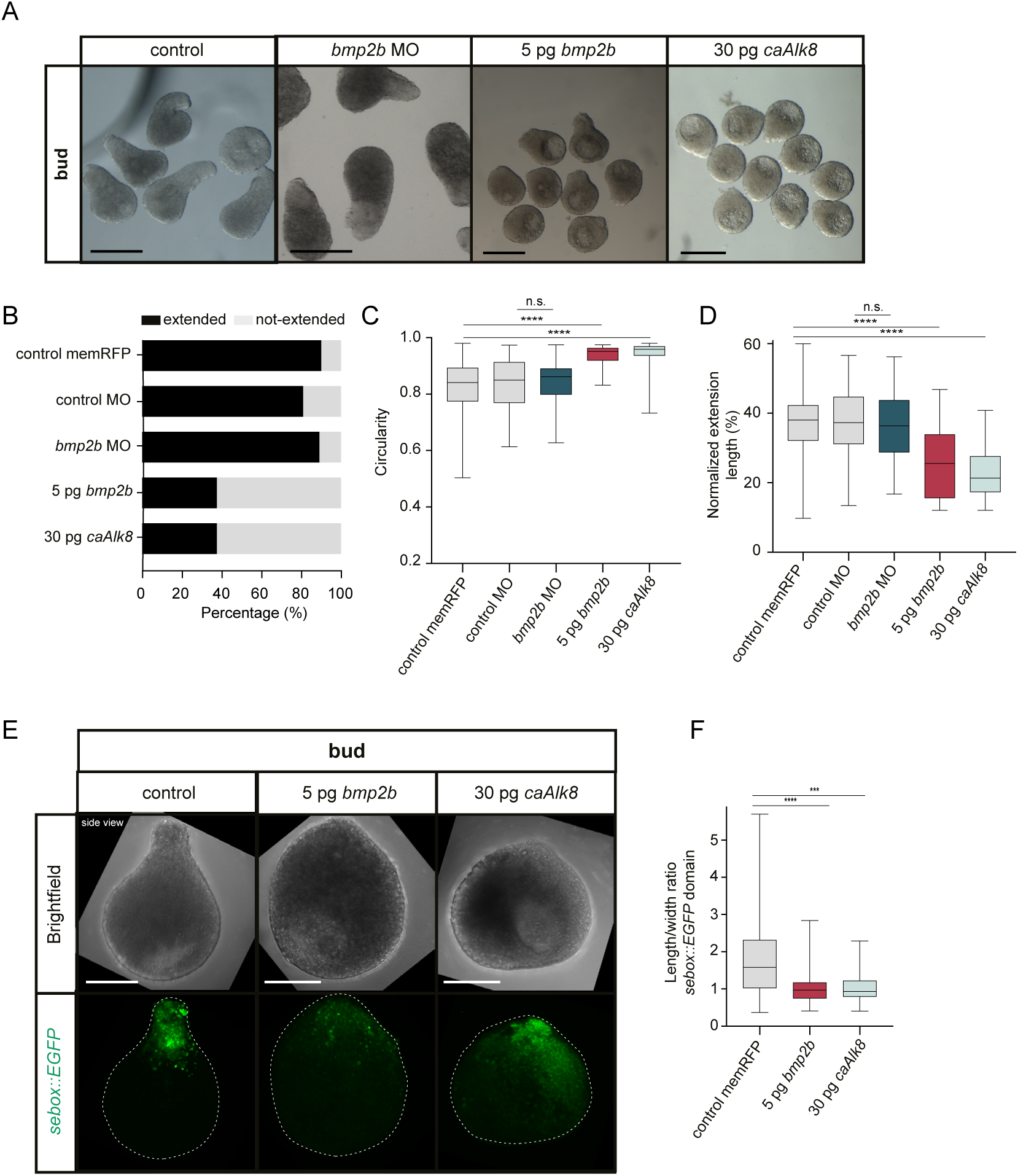
Changes in blastoderm explant elongation upon BMP signaling overactivation. (**A**) Single-plane bright-field images of blastoderm explants from wildtype embryos (control: n=191, N=11), and embryos injected with 1.5ng *bmp2b morpholino* (MO) (n=55, N=5), 5pg *bmp2b* (n=56, N=5) or 30pg *caAlk8* mRNA (n=112, N=6) (side views) at bud stage. All embryos were co-injected with 50-100pg *memRFP* or *memGFP* as injection control. (**B**) Percentage of extended or not-extended blastoderm explants from wildtype embryos (control: n=191, N=11; 5ng controlMO: n=63, N=5), and embryos injected with 1.5ng *bmp2b*MO (n=55, N=5), 5pg *bmp2b* (n=56, N=5) or 30pg *caAlk8* mRNA (n=112, N=6) at bud stage. All embryos were co-injected with 50-100pg *memRFP* or *memGFP* as injection control. (**C**) Circularity of extended or not-extended blastoderm explants from wildtype embryos (control: n=191, N=11; controlMO: n=63, N=5), and embryos injected with 1.5ng *bmp2b*MO (n=55, N=5), 5pg *bmp2b* (n=56, N=5) or 30pg *caAlk8* mRNA (n=112, N=6) at bud stage. All embryos were co-injected with 50-100pg *memRFP* or *memGFP* as injection control. ****p<0.0001, ns, not significant (Kruskal-Wallis test). (**D**) Normalized extension length of blastoderm explants from wildtype embryos (control: n=172, N=11; controlMO: n=51, N=5), and embryos injected with 1.5ng *bmp2b*MO (n=49, N=5), 5pg *bmp2b* (n=21, N=5) or 30pg *caAlk8* mRNA (n=45, N=6) at bud stage. All embryos were co-injected with 50-100pg *memRFP* or *memGFP* as injection control. ****p<0.0001, ns, not significant (One-way ANOVA). (**E**) Maximum intensity projection of brightfield/fluorescence images (side views) of blastoderm explants from Tg(*sebox::EGFP*) wildtype embryos expressing EGFP (green) in mesendoderm progenitors (control: n=62, N=8), and Tg(*sebox::EGFP*) embryos injected with 5pg *bmp2b* (n=33, N=6) or 30pg *caAlk8* mRNA (n=31, N=4) at bud stage. All embryos were co-injected with 50-100pg *memRFP* as injection control. (**F**) Length/width ratio of the EGFP expression domain in blastoderm explants from Tg(*sebox::EGFP*) wildtype embryos expressing EGFP (green) in mesendoderm progenitors (control: n=62, N=8), and Tg(*sebox::EGFP*) embryos injected with 5pg *bmp2b* (n=33, N=6) or 30pg *caAlk8* mRNA (n=31, N=4) at bud stage. All embryos were co-injected with 50-100pg *memRFP* as injection control. ****p<0.0001, ***p=0.0002 (Kruskal-Wallis test). Scale bars: 500 µm (A), 200 µm (E).

To further challenge this notion, we locally overactivated BMP signaling by creating large cell clones overexpressing *caAlk8* within the explants. We found that explants expressing *caAlk8* predominantly in mesendodermal progenitors exhibited the most severe elongation defects (Fig. S3G-I), suggesting that the observed explant elongation defects in response to increased BMP signaling were largely due to excessive BMP signaling within the mesendoderm.

### BMP signaling reduces mesendodermal cell dispersal and cell alignment

Next, we asked how increased BMP signaling within the mesendoderm reduces explant elongation. Given that BMP signaling has been suggested to spatially restrict oriented cell intercalations to the dorsal side in the zebrafish gastrula (Myers *et al*, 2002a), we analyzed whether overactivation of BMP in the mesendoderm might reduce explant elongation by inhibiting cell intercalation within the explant extension. Comparing the behavior of clones in the mesendoderm in control explants to *caAlk8* overexpressing explants with elevated BMP signaling levels revealed that clone elongation along the direction of the extension was reduced upon BMP overactivation (2.02-fold mean increase in clone length in control conditions versus 1.28-fold increase in clone length in *caAlk8* explants) (Fig. 3A,B, Fig. S4A, Supplementary Video 5). In line with this, cells in *caAlk8* overexpressing explants failed to effectively polarize and align along the mediolateral explant axis during explant elongation (Fig. 3C,D), indicative of reduced medially-oriented cell intercalations. Other functions of BMP signaling in mesendoderm morphogenesis, such as regulating the direction of mesendoderm migration during convergence movements (von der Hardt *et al*, 2007), were unlikely to represent main effector processes by which excessive BMP signaling interferes with explant elongation, as decreasing BMP signaling activity (Fig. 2A-D, Fig. S2D-G) or knocking down the expression of *has2* (Fig. S4B-D), a critical regulator of convergence movements in the embryo (Bakkers *et al*, 2004), had no clear effect on explant elongation.

**Figure 3.**
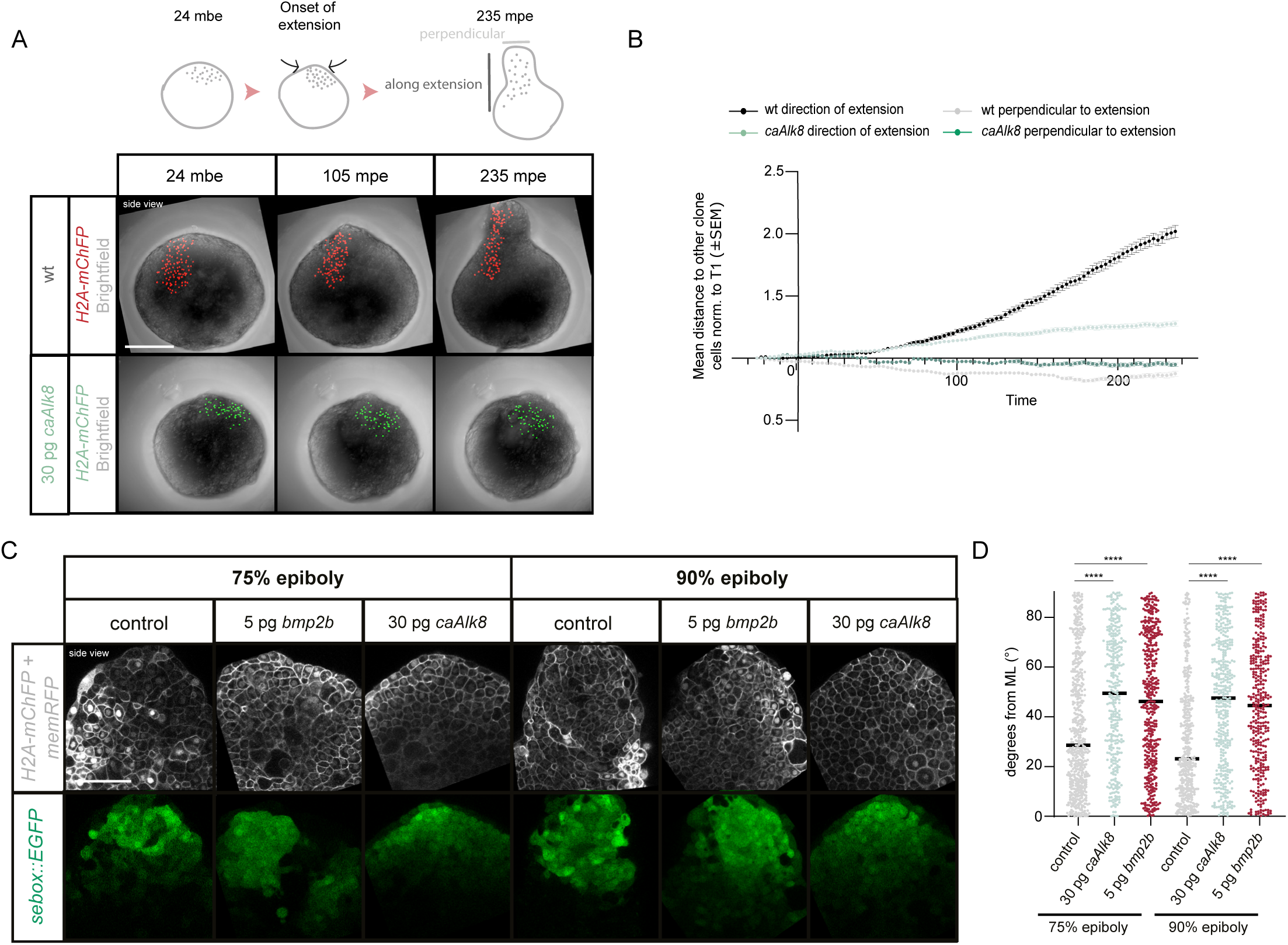
Changes in clone elongation and cell alignment in blastoderm explants upon BMP signaling overactivation. (**A**) Maximum intensity projection of fluorescence/brightfield images (side views) of blastoderm explants before explant elongation (24 mbe, left column), during explant elongation (105 mpe, middle column) and at the end of elongation (235 mpe, right column) showing clonally labeled cell nuclei in the mesendoderm in blastoderm explants from wildtype embryos (red) or embryos injected with 30pg *caAlk8* mRNA (green). Images for wildtype explants correspond to Fig. 1G. (**B**) Mesendodermal clone dispersal parallel and perpendicular to the axis of explant elongation assessed by the mean distance of each cell in the clone to other clone cells during explant elongation in blastoderm explants from wildtype embryos (black/grey) or embryos injected with 30pg *caAlk8* mRNA (light/dark green) (wildtype >320 cells analyzed per time point, n=5, N=5; 30pg *caAlk8* >420 cells analyzed per time point, n=5, N=5). Data for wildtype explants corresponds to Fig. 1H. (**C**) Single-plane high-resolution images (side views) of blastoderm explants obtained from Tg(*sebox::EGFP*) embryos expressing EGFP (green) in mesendoderm progenitors, and Tg(*sebox::EGFP*) embryos injected with 5pg *bmp2b* and 30pg *caAlk8* during explant elongation (corresponding to embryonic 75% and 90% epiboly stage respectively). Cell outlines are marked by 80-100pg *memRFP* mRNA injection (grey). Cell nuclei are marked by 40pg *H2A-mChFP* mRNA injection (grey). (**D**) Cell alignment assessed by the deviation (degrees) of the main cell extension axis from the main mediolateral (ML) explant axis during explant elongation (75% epiboly: wildtype: 545 cells, n=12, N=6; 5pg *bmp2b*: 417 cells, n=11, N=6; 30pg *caAlk8*: 342 cells, n=11, N=6; 90% epiboly: wildtype: 439 cells, n=12, N=7; 5pg *bmp2b*: 347 cells, n=11, N=7; 30pg *caAlk8*: 399 cells, n=11, N=7). ****p<0.0001 (Kruskal-Wallis test). Scale bars: 200 µm (A), 100 µm (C).

Collectively, these observations suggest that mesendoderm ventralization by BMP overactivation blocks explant elongation by reducing medially-oriented mesendodermal cell alignment and intercalation.

### BMP signaling is attenuated in the mesendoderm by Nodal signaling activity

Given that spatially confining BMP signaling activity is important for explant elongation, we next asked how BMP signaling activity is distributed within the explants. In line with previous observations (Fulton *et al*, 2020; Schauer *et al*, 2020), we found that a long-range gradient of BMP signaling, as monitored by the nuclear localization of the phosphorylated and thus activated form of the BMP downstream signaling mediator and transcriptional regulator Smad5 (pSmad5), was formed across the explant during the elongation phase with the lowest pSmad5 levels overlapping with the site of elongation (Fig. 4A,B). Interestingly, when we analyzed the pSmad5 activity domain in a *sebox::EGFP* background, we noted that the mesendodermal *sebox::EGFP*-positive domain was largely devoid of pSmad5, suggesting that BMP signaling was inherently suppressed in the mesendoderm of explants (Fig. 4A,B).

**Figure 4.**
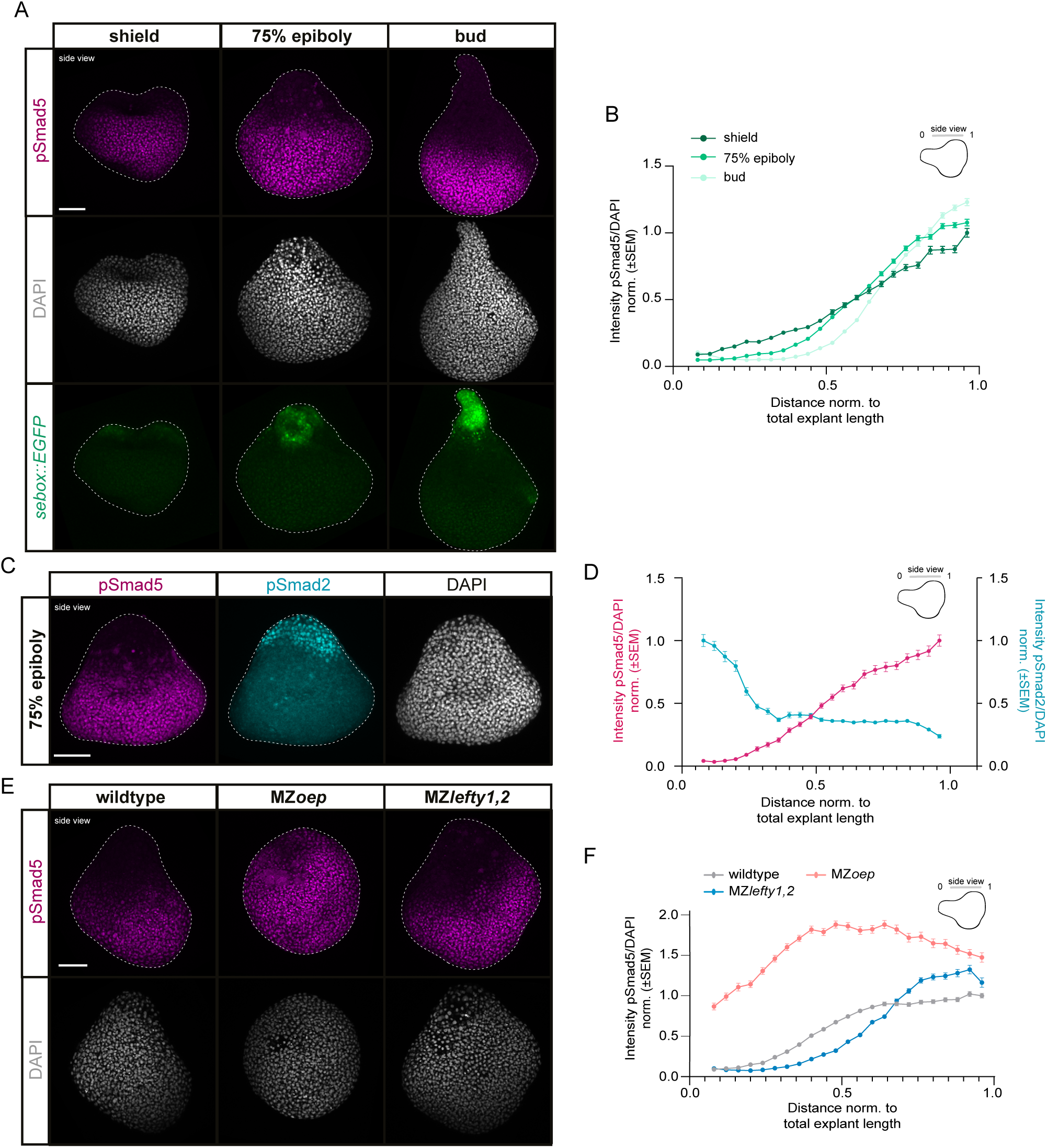
Repression of BMP signaling activity by Nodal signaling at the sites of mesendoderm induction and blastoderm explant elongation. (**A**) Maximum intensity projection of fluorescence images (side views) of blastoderm explants from Tg(*sebox::EGFP*) embryos marking mesendoderm progenitors (green) during explant elongation (corresponding to embryonic shield: n=6, N=3; 75% epiboly: n=6, N=3; bud stage: n=6, N=3) stained for pSmad5 (BMP signaling activity; magenta) and DAPI (nuclei; grey). (**B**) Intensity of nuclear pSmad5 normalized to DAPI as a function of the distance from the explant tip during explant elongation (corresponding to embryonic shield: n=6, N=3; 75% epiboly: n=6, N=3; bud stage: n=6, N=3). Intensities are shown relative to the mean intensity in the bin closest to the back (high intensity domain) of shield stage explants. Notably the first and last 4% of the explant were excluded due to low number of nuclei at the sample edges. (**C**) Maximum intensity projection of fluorescence images (side views) of blastoderm explants during explant elongation (corresponding to embryonic 75% epiboly stage) stained for pSmad5 (BMP signaling activity; magenta), pSmad2 (Nodal signaling activity; cyan) and DAPI (nuclei; grey) (n=6, N=3). (**D**) Intensity of nuclear pSmad5 (BMP signaling activity; magenta) and pSmad2 (Nodal signaling activity; cyan) normalized to DAPI as a function of the distance from the explant tip during explant elongation (corresponding to embryonic 75% epiboly stage) (n=6, N=3). Intensities are shown relative to the mean intensity in the bin closest to the back (high intensity domain) of the explants for pSmad5 and closest to the tip of the explant for pSmad2 (high intensity domain). Notably the first and last 4% of the explant were excluded due to low number of nuclei at the sample edges. (**E**) Maximum intensity projection of fluorescence images (side views) of blastoderm explants from wildtype, MZ*oep* and MZ*lefty1,2* embryos during explant elongation (corresponding to embryonic 75% epiboly stage) stained for pSmad5 (BMP signaling activity; magenta) and DAPI (nuclei; grey) (wildtype: n=9, N=4; MZ*oep*: n=9, N=4; MZ*lefty1,2*: n=9, N=4). (**F**) Intensity of nuclear pSmad5 normalized to DAPI as a function of the distance from the explant tip during elongation (corresponding to embryonic 75% epiboly stage) (wildtype: n=9, N=4; MZ*oep*: n=9, N=4; MZ*lefty1,2*: n=9, N=4). Intensities are shown relative to the mean intensity in the first bin closest to the back (high intensity domain) of the wildtype explants. Notably the first and last 4% of the explant were excluded due to low number of nuclei at the sample edges. Scale bars: 100 µm (A,C,E).

As mesendoderm induction depends on Nodal signaling (Schier & Talbot, 2005; Schier, 2009; Rogers & Müller, 2019; Economou & Hill, 2020; Hill, 2022), we asked whether Nodal signaling might also negatively regulate the extent of BMP signaling concomitantly with its activity in inducing mesendodermal cell fates. Consistent with such a role of Nodal signaling, the expression of several components of the BMP signaling pathway has previously been shown to be affected by changes in Nodal signaling activity in intact embryos (Gritsman *et al*, 1999; Sirotkin *et al*, 2000; Lele *et al*, 2001b; Bennett *et al*, 2007; Varga *et al*, 2007; Xu *et al*, 2014; Wang *et al*, 2021; Cheng *et al*, 2023). Such a function of Nodal signaling is likely to be restricted to later gastrulation stages, as at the onset of gastrulation, the loss of Nodal signaling activity in intact embryos does not obviously affect overall pSmad5 gradient formation (Rogers *et al*, 2020). Supporting a role of Nodal signaling in restricting BMP signaling within the explant, we found that the activities of pSmad5 and the Nodal downstream signaling mediator and transcriptional regulator pSmad2 peaked in opposing domains during explant elongation (Fig. 4C,D). Moreover, in explants prepared from mutant embryos with increased Nodal signaling activity (MZ*lefty1,2*) (Rogers *et al*, 2017), or devoid of active Nodal signaling (MZ*oep*) (Gritsman *et al*, 1999), the extent of the domain of high nuclear pSmad5 was reduced (MZ*lefty1,2*) or clearly expanded (MZ*oep*) along the back-tip explant axis (Fig. 4E,F, Fig. S5A-E), demonstrating a critical role of Nodal signaling in restricting BMP from the mesendodermal portion of the elongating explant. Given that wildtype and MZ*lefty1,2* explants were similar in length at 75% epiboly (Fig. S5F), and that there was no clear correlation between explant length and pSmad5 domain extent along the explant axis (Fig. S5G), this activity of Nodal signaling in restricting BMP signaling appears to be due to a signaling function of Nodal, changing also the amount of mesendoderm (Fig. S5H), rather than being the secondary consequence of Nodal-induced morphogenesis, previously shown to separate the domains of BMP and Wnt signaling activity for neuroectoderm patterning (Fulton *et al*, 2020).

### Nodal represses BMP signaling activity during explant elongation by upregulation of *chrd* expression

Next, we asked how Nodal signaling couples mesendoderm induction with the restriction of BMP signaling activity, two prerequisites for explant elongation (Fulton *et al*, 2020; Schauer *et al*, 2020; Williams & Solnica-Krezel, 2020b) (Fig. 1C-H, Fig. S1F,F’, Fig.2A-F). Ubiquitous overexpression of the Nodal ligand *cyclops* completely blocked BMP signaling activity as monitored by the lack of nuclear pSmad5 localization in embryos consisting of only mesendoderm (Fig. S6A,B). However, co-injecting the constitutively active BMP receptor *caAlk8* could effectively recover BMP signaling activity in Nodal overexpressing embryos (Fig. S6A,B), suggesting that Nodal signaling antagonizes BMP signaling activity predominantly by regulation of extracellular or surface-bound BMP signaling effectors.

The expression domain of one of the main BMP antagonists, the dorsally-expressed *chrd* (Hammerschmidt *et al*, 1996; Schulte-Merker *et al*, 1997; Miller-Bertoglio *et al*, 1997), has previously been shown to be modulated by Nodal signaling (Gritsman *et al*, 1999; Sirotkin *et al*, 2000; Bennett *et al*, 2007; Varga *et al*, 2007; Xu *et al*, 2014; Cheng *et al*, 2023). Hence, we sought to assess a potential requirement of Nodal-dependent *chrd* activation for restricting BMP signaling at the site of explant elongation. To this end, we first analyzed changes in *chrd* expression in explants obtained from MZ*lefty1,2* (Rogers *et al*, 2017) and MZ*oep* (Gritsman *et al*, 1999) mutant embryos, where Nodal signaling is up- and down-regulated, respectively, during explant elongation, i.e. corresponding to embryonic 75% epiboly stage. While increasing Nodal signaling above wildtype levels in MZ*lefty1,2* explants led to mildly elevated levels of *chrd* expression (Fig. 5A-D; note that the upregulation of *chrd* expression in MZ*lefty1,2* explants was variable), reduced Nodal signaling in MZ*oep* mutant explants strongly diminished *chrd* expression during explant elongation as shown by *in situ* hybridization and qPCR analysis (Fig. 5A-D). This suggests that the activation of Nodal signaling is required for properly establishing the *chrd* expression domain within explants.

**Figure 5.**
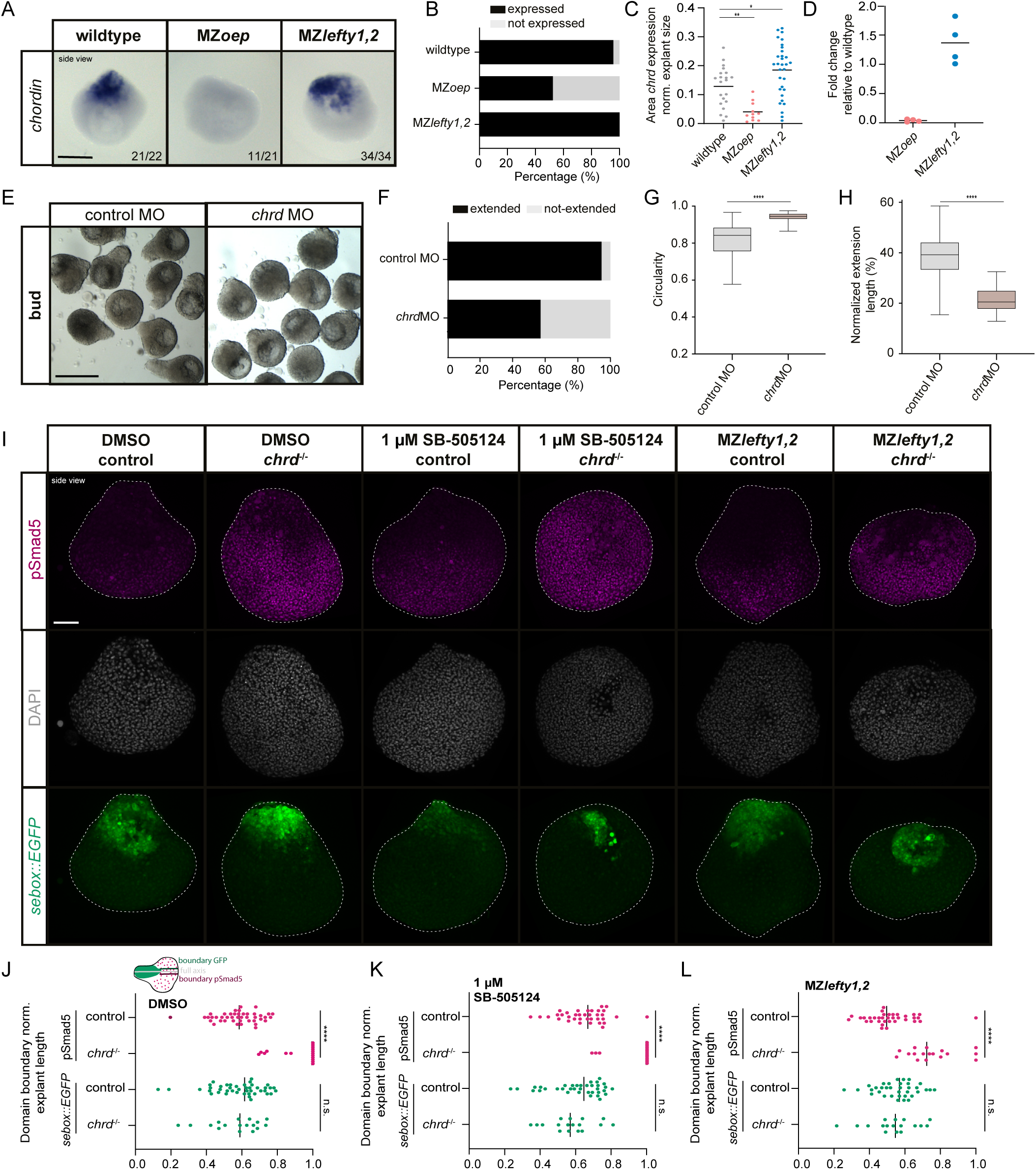
Restriction of BMP signaling activity and explant elongation by Nodal-dependent regulation of *chordin*. (**A**) Expression of *chordin* (*chrd*) assessed by *in situ* hybridization in blastoderm explants from wildtype, MZ*oep* and MZ*lefty1,2* embryos (side view) during explant elongation (corresponding to embryonic 75% epiboly stage). The proportion of explants with a similar expression pattern to the shown image is shown in the lower right corner (wildtype: n=22, N=3; MZ*oep*: n=21, N=3; *MZlefty1,2*: n=34, N=3). (**B**) Percentage of expressing/not expressing blastoderm explants from wildtype (n=23, N=3), MZ*oep* (n=21, N=3) and MZ*lefty1,2* (n=34, N=3) embryos corresponding to 75% epiboly stage. (**C**) Area of the expression domain of *chrd* assessed by *in situ* hybridization normalized to explant area in blastoderm explants from wildtype (n=21, N=3), MZ*oep* (n=10, N=3) and MZ*lefty1,2* (n=34, N=3) embryos during explant elongation (corresponding to embryonic 75% epiboly stage). The black line indicates the mean. *p=0.0229, **p=0.0061 (One-way ANOVA). (**D**) Fold change of *chrd* expression in blastoderm explants from MZ*oep* (N=4) and MZ*lefty1,2* (N=4) embryos relative to blastoderm explants from wildtype embryos during explant elongation (corresponding to embryonic 75% epiboly stage). The black line indicates the mean. (**E**) Single-plane bright-field images (side views) of blastoderm explants obtained from wildtype embryos (control MO: n=74, N=4) and embryos injected with 4.5ng *chrd*MO (n=56, N=5) at bud stage. All embryos were co-injected with 50-100pg *memRFP* or *memGFP* as injection control. (**F**) Percentage of extended/not-extended blastoderm explants from wildtype embryos (control MO: n=74, N=4) and embryos injected with 4.5ng *chrd*MO (n=56, N=5) at bud stage. All embryos were co-injected with 50-100pg *memRFP* or *memGFP* as injection control. (**G**) Circularity of extended/not-extended blastoderm explants from wildtype embryos (control MO: n=74, N=4) and embryos injected with 4.5ng *chrd*MO (n=56, N=5) at bud stage. All embryos were co-injected with 50-100pg *memRFP* or *memGFP* as injection control. ****p<0.0001 (Mann-Whitney test). (**H**) Normalized extension length of extended blastoderm explants from wildtype embryos (control MO: n=70, N=4) and embryos injected with 4.5ng *chrd*MO (n=28, N=5) at bud stage. All embryos were co-injected with 50-100pg *memRFP* or *memGFP* as injection control. ****p<0.0001 (Unpaired t-test). (**I**) Maximum intensity projection of fluorescence images (side views) of blastoderm explants from wildtype embryos and blastoderm explants from *chrd*^-/-^ mutant embryos treated with DMSO (treated from 256c to 75% epiboly: wildtype: n=43, N=6; *chrd*^-/-^: n=17, N=4) or 1 µM Nodal inhibitor (SB-505124; treated from 256c to 75% epiboly: wildtype: n=36, N=6; *chrd*^-/-^: n=17, N=6) and blastoderm explants from MZ*lefty1,2* (n=35, N=5) and MZ*lefty1,2;chrd*^-/-^ (n=17, N=4) mutant embryos with Tg(*sebox::EGFP*) marking mesendoderm progenitors (green) during explant elongation (corresponding to embryonic 75% epiboly stage) stained for pSmad5 (BMP signaling activity; magenta) and DAPI (nuclei; grey). (**J-L**) Domain of nuclear pSmad5 and EGFP expression measured centrally from the back of the explant normalized to the explant length in blastoderm explants obtained from wildtype embryos and from *chrd*^-/-^ embryos treated with (J) DMSO (treated from 256c to 75% epiboly: wildtype: n=43,N=6; *chrd*^-/-^: n=17, N=4) or (K) 1 µM Nodal inhibitor (SB-505124; treated from 256c to 75% epiboly: wildtype: n=36, N=6; *chrd*^-/-^: n=17, N=6) and from (L) MZ*lefty1,2* (n=35, N=5) and MZ*lefty1,2;chrd*^-/-^ (n=17, N=4) mutant embryos during explant elongation (corresponding to embryonic 75% epiboly stage) with Tg(*sebox::EGFP*) marking mesendoderm progenitors. ****p<0.0001, ns, not significant (Mann-Whitney test: (J) *sebox::EGFP* domain extent and (K, L) pSmad5 domain extent; Unpaired t-test: (J) pSmad domain extent and (K, L) *sebox::EGFP* domain extent). Scale bars: 200 µm (A), 500 µm (E), 100 µm (I).

To assess the requirement of *chrd*-mediated BMP signaling restriction for explant elongation, we analyzed how *chrd* loss-of-function affects explant elongation. To this end, we generated blastoderm explants from *chrd* morphant embryos (Nasevicius & Ekker, 2000). These *chrd* morphant explants showed a reduction in explant elongation frequency and length (Fig. 5E-H) and a reduced length/width ratio of the mesendoderm (Fig. S6C,D), reminiscent of explants overexpressing *bmp* ligands or *caAlk8*. This suggests that proper *chrd* function is critical for allowing effective explant elongation.

To determine whether *chrd* is required for restricting BMP signaling activity downstream of Nodal, we analyzed the ability of Nodal signaling to modulate BMP signaling, as monitored by nuclear pSmad5 localization, in explants obtained from *chrd* mutant embryos (Hammerschmidt *et al*, 1996). To this end, we prepared explants from wildtype and *chrd* mutant embryos (Hammerschmidt *et al*, 1996) under normal Nodal signaling conditions (wildtype or *chrd*^-/-^ mutant explants exposed to DMSO), reduced Nodal signaling conditions (wildtype or *chrd*^-/-^ mutant explants exposed to 1 µM of the Nodal signaling inhibitor SB-505124 (Byfield *et al*, 2004; Rogers *et al*, 2017)) or increased Nodal signaling (MZ*lefty1,2* (Rogers *et al*, 2017) or MZ*lefty1,2;chrd*^-/-^ triple mutant explants) (Fig. 5I-L). While the relative positioning of the *sebox::EGFP* expression domain along the main explant axis seemed largely unaffected in *chrd*^-/-^ explants (Fig. 5I-L), the pSmad5 activity domain was strongly expanded into the mesendoderm irrespective of the state of Nodal signaling within the explant (Fig. 5I-L, S6E,F). Notably, an area devoid of nuclear pSmad5 was still formed in some *chrd*^-/-^ explants, in particular when Nodal signaling levels were increased (MZ*lefty1,2;chrd*^-/-^ explants) (Fig. 5I-L, S6E,F), suggesting that additional *chrd*-independent mechanisms might be activated by peak Nodal signaling levels to further repress BMP signaling. Overall, this suggests that Nodal signaling attenuated BMP signaling within the mesendoderm by activating *chrd* expression in explants, with additional *chrd*-independent mechanisms maintaining low levels of pSmad5 in the mesendoderm under high Nodal signaling conditions. Thus, Nodal signaling drives explant elongation not only by inducing mesendoderm, but also by restricting BMP signaling activity via *chordin*-dependent and *chordin*-independent mechanisms at the site of explant extension.

### Nodal signaling maintains an area of low BMP signaling on the dorsal side of the gastrula for robust axis elongation

Given the key role of Nodal signaling in mediating explant elongation by both inducing mesendoderm specification and restricting BMP expression, we asked whether a similar function for Nodal signaling exists in the embryo. To this end, we first tested whether Nodal signaling can also modulate the long-range pSmad5 gradient, extending from ventral to the dorsal side in the embryo (Ramel & Hill, 2013; Tucker *et al*, 2008; Zinski *et al*, 2017; Pomreinke *et al*, 2017; Rogers *et al*, 2020; Greenfeld *et al*, 2021), as found for explants (Fig. 4E,F). In mutant embryos with reduced (MZ*oep*) or increased (MZ*lefty1,2*) Nodal signaling levels, the BMP signaling range, as monitored by nuclear pSmad5 localization, was expanded or reduced, respectively at 75% epiboly (Fig. 6A,B, S7A-D’’’), thereby mimicking the situation in explants. Specifically, we found that MZ*oep* mutant embryos at 75% epiboly showed an overall increase of high nuclear pSmad5 levels compared to wildtype embryos at the same stage (Fig. 6A,B, S7A,B,D-D’’’), while the domain of high pSmad5 signaling activity was more restricted towards the ventral side in MZ*lefty1,2* mutant embryos (Fig. 6A,B, S7A,C,D-D’’’). Collectively, this suggests that Nodal signaling in embryos at mid-gastrula stages displayed a similar, but also less pronounced ability in restricting high BMP signaling to the ventral half of the embryo as found in the mesendoderm of explants.

**Figure 6.**
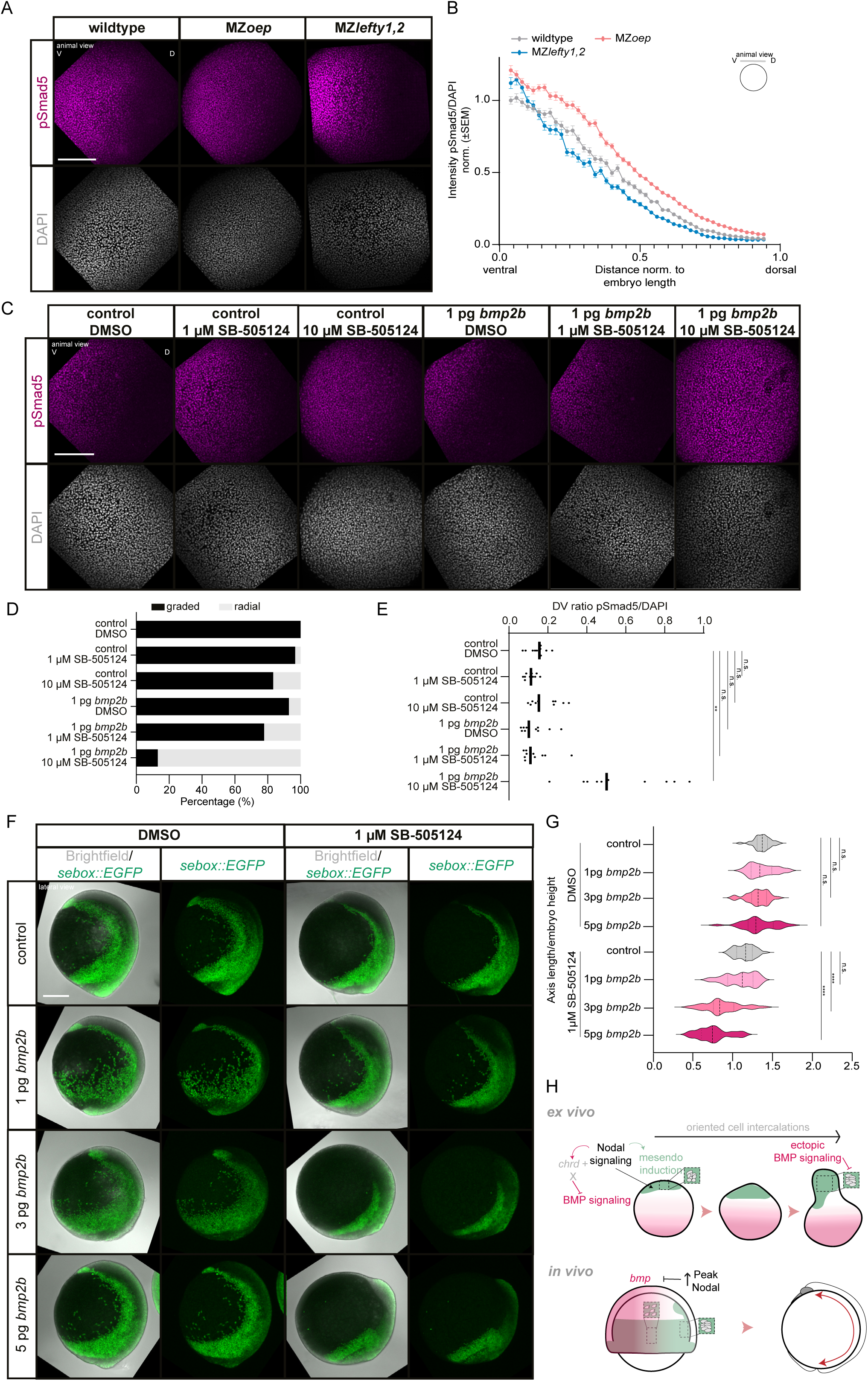
Suppression of BMP signaling on the dorsal side by peak Nodal signaling levels providing robustness for embryonic body axis elongation. (**A**) Maximum intensity projection of fluorescence images (animal views) of wildtype, MZ*oep* and MZ*lefty1,2* embryos at 75% epiboly stage stained for pSmad5 (BMP signaling activity; magenta) and DAPI (nuclei; grey) (wildtype: n=11, N=5; MZ*oep*: n=11, N=5; MZ*lefty1,2*: n=11, N=5). D=dorsal, V=ventral. (**B**) Intensity of nuclear pSmad5 normalized to DAPI as a function of the distance from the ventral side at 75% epiboly stage (wildtype: n=11, N=5; MZ*oep*: n=11, N=5; MZ*lefty1,2*: n=11, N=5). Intensities are shown relative to the mean intensity in the first bin closest to the ventral side (high intensity domain) of wildtype embryos. Notably the first and last 4% of the embryo were excluded due to low number of nuclei at the sample edges. (**C**) Maximum intensity projection of fluorescence images (animal views) of DMSO treated (4-16c>75% epiboly, control: n=35, N=6; 1pg *bmp2b*: n=28, N=6), 1 µM SB-505124 treated (4-16c>75% epiboly, control: n=30, N=6; 1pg *bmp2b*: n=27, N=6) and 10 µM SB-505124 treated (4-16c>75% epiboly, control: n=36, N=6; 1pg *bmp2b*: n=23, N=6) embryos at 75% epiboly stage stained for pSmad5 (BMP signaling activity; magenta) and DAP (nuclei; grey). All embryos were co-injected with 80pg *memGFP* as injection control. (**D**) Proportion of DMSO treated (4-16c>75% epiboly, control: n=35, N=6; 1pg *bmp2b*: n=28, N=6), 1 µM SB-505124 treated (4-16c>75% epiboly, control: n=30, N=6; 1pg *bmp2b*: n=27, N=6) and 10 µM SB-505124 treated (4- 16c>75% epiboly, control: n=36, N=6; 1pg *bmp2b*: n=23, N=6) embryos at 75% epiboly stage exhibiting a gradient of nuclear pSmad5 or a radial pSmad5 domain. All embryos were co-injected with 80pg *memGFP* as injection control. (**E**) Ratio of dorsal pSmad5 intensities relative to ventral pSmad5 intensities normalized to DAPI of DMSO treated (4-16c>75% epiboly, control: n=11, N=5; 1pg *bmp2b*: n=11, N=5), 1 µM SB-505124 treated (4-16c>75% epiboly, control: n=11, N=5; 1pg *bmp2b*: n=11, N=5) and 10 µM SB-505124 treated (4-16c>75% epiboly, control: n=11, N=5; 1pg *bmp2b*: n=11, N=5) embryos at 75% epiboly stage. All embryos were co-injected with 80pg *memGFP* as injection control. **p=0.0045, ns, not significant (Kruskal-Wallis test) (**F**) Maximum intensity projection of brightfield/fluorescence images (lateral views) of DMSO treated (4-16c>75% epiboly, control: n=30, N=7; 1pg *bmp2b*: n=28,N=7; 3pg *bmp2b*: n=22, N=7; 5pg *bmp2b*: n=25,N=7) (left side) or 1 µM SB-505124 treated (4-16c>75% epiboly, control: n=40, N=7; 1pg *bmp2b*: n=31, N=7; 3pg *bmp2b*: n=28, N=7; 5pg *bmp2b*: n=28, N=7) (right side) embryos at bud stage expressing Tg(*sebox::EGFP*) to mark mesendodermal progenitors (green). All embryos were co-injected with 80pg *memRFP* as injection control. (**G**) Ratio of mesendodermal AP axis length/embryo height of DMSO treated (4-16c>75% epiboly, control: n=30, N=7; 1pg *bmp2b*: n=28, N=7; 3pg *bmp2b*: n=22, N=7; 5pg *bmp2b*: n=25, N=7) or 1 µM SB-505124 treated (4-16c>75% epiboly, control: n=40, N=7; 1pg *bmp2b*: n=31, N=7; 3pg *bmp2b*: n=28, N=7; 5pg *bmp2b*: n=28, N=7) embryos at bud stage. All embryos were co-injected with 80pg *memRFP* as injection control. ****p<0.0001 (One-way ANOVA). (**H**) Schematic of Nodal-dependent restriction of BMP signaling activity to allow effective axial elongation by cell intercalations *ex vivo* and *in vivo*. Scale bars: 200 µm (A,C,F).

To further challenge the hypothesis that Nodal signaling maintains an area of low BMP signaling at the dorsal side of the embryo, we analyzed changes in nuclear pSmad5 levels on the dorsal side of the embryo upon *bmp2b* overexpression under normal and reduced Nodal signaling conditions. To this end, we injected low amounts of *bmp2b* mRNA, which on its own would not abolish pSmad5 gradient formation (Fig. 6C,D), in embryos exposed to 1 µM or 10 µM of the Nodal signaling inhibitor SB-505124 (Byfield *et al*, 2004). We found that with increasing concentration of the Nodal inhibitor added, embryos became more sensitive to the *bmp2b* overexpression, changing the pSmad5 gradient profile and leading to embryos displaying near-uniform high levels of nuclear pSmad5 upon simultaneous BMP overexpression and Nodal signaling reduction (Fig. 6C-E). This showed that Nodal signaling at the dorsal side of the embryo is required to effectively repress BMP signaling, like in the explant mesendoderm.

When analyzing the effect of BMP overactivation on axis elongation in intact embryos by injecting 30pg *caAlk8*, we found that the AP length of the mesendodermal domain was reduced by the end of gastrulation (Fig.S7E,F), consistent with previous findings (Myers *et al*, 2002a), and reminiscent of our observations made in explants (Fig. 2). Furthermore, we observed that cell alignment along the mediolateral axis within the dorsal mesendoderm was disrupted upon BMP overactivation at late gastrula (90% epiboly) (Fig. S7G,H), suggesting that, similar to the situation in explants (Fig. 3), excessive BMP signaling reduces effective mesendoderm C&E movements by reducing medially-oriented mesendoderm cell alignment and thus intercalations.

Finally, to test the effect of the interaction of BMP and Nodal signaling on axis elongation in the embryo, we asked whether peak levels of Nodal signaling also provide robustness for embryo body axis extension by reducing the sensitivity towards BMP signaling in the dorsal domain. In embryos injected with various low amounts of *bmp2b* mRNA (1-5pg), body axis extension remained largely unchanged as compared to control embryos (Fig. 6F,G). However, when we simultaneously treated BMP overexpressing embryos with 1 µM SB-505124 to reduce peak Nodal signaling levels, we found that axis elongation was severely reduced (Fig. 6F,G). This points to a conserved function of Nodal signaling in axis extension by effectively repressing BMP signaling at the dorsal side of both embryos and explants.

## Discussion

Our study describes a so far understudied role of Nodal signaling in rendering the dorsal mesendoderm less sensitive to BMP signaling during gastrulation movements, thereby providing robustness to axis elongation. Robustness towards extrinsic and intrinsic perturbations is an important feature for development to have a reproducible outcome (Wolpert, 1992; Kitano, 2004; Masel & Siegal, 2009) with mechanochemical feedback loops, linking signaling, patterning and morphogenesis, constituting key mechanisms securing robustness (Gilmour *et al*, 2017; Hannezo & Heisenberg, 2019; Collinet & Lecuit, 2021; Bailles *et al*, 2022). In the present study, we identify a critical role for morphogen signaling pathway crosstalk in maintaining the relative spatial organization of distinct signaling domains within the gastrula (Fig. 6H), a prerequisite for embryo axis elongation to occur normally even upon variations in morphogen signaling and the overall embryonic context. Cross-regulation between BMP and Nodal signaling displaying opposing morphogenetic activities has also been implicated in the emergence of left-right heart asymmetry in zebrafish development (Veerkamp *et al*, 2013), although how such coordination is realized in different geometric and signaling contexts has not been addressed yet.

Nodal and BMP signaling form activity gradients along the marginal-to-animal and ventral-to-dorsal axes of the zebrafish gastrula, respectively (Schier & Talbot, 2005; Rogers & Müller, 2019; Hill, 2022), which only partially overlap, a spatial arrangement critical for proper embryonic axis formation and patterning (Fauny *et al*, 2009; Xu *et al*, 2014; Soh *et al*, 2020). Consistent with their partially distinct activity domains, Nodal and BMP signaling can exert opposite functions during patterning by e.g. inducing dorsally-derived head structures (Gritsman *et al*, 2000) and ventral tissues (Kishimoto *et al*, 1997; Nguyen *et al*, 1998; Hild *et al*, 1999; Dick *et al*, 2000; Schmid *et al*, 2000), respectively, and during morphogenesis by e.g. differentially regulating cell-cell adhesion (von der Hardt *et al*, 2007; Barone *et al*, 2017). Hence, the effective coordination of the domains of Nodal and BMP signaling activity is essential to not only properly pattern, but also shape the gastrula stage embryo, consistent with our observation that ectopic BMP signaling in the dorsal mesendoderm disrupts Nodal-induced cell intercalations and thus axis elongation.

Beyond this necessity to spatially coordinate Nodal and BMP signaling activity for axis elongation, our results show that Nodal signaling actually represses BMP signaling thereby creating an area of low sensitivity towards BMP at the site of Nodal-mediated mesendoderm induction. Interestingly, although Nodal signaling is required for the early expression of BMP regulators, such as *chrd* or *noggin1* together with β-catenin signaling on the dorsal side of the gastrula (Gritsman *et al*, 1999; Sirotkin *et al*, 2000; Varga *et al*, 2007), the formation of the pSmad5 gradient has been shown to be unaffected by loss of Nodal signaling activity at the onset of gastrulation (Rogers *et al*, 2020). This lack of repressing activity of Nodal signaling on BMP activity might be restricted to pre-gastrula stages since increased Nodal signaling as observed in Nodal type II receptor loss-of-function is accompanied by a decrease of BMP signaling activity during gastrulation (Preiß *et al*, 2022). Our data show that indeed at late gastrulation stages, when cells undergo large-scale cell rearrangements associated with C&E movements, peak levels of Nodal signaling in the dorsal mesendoderm render the tissue less sensitive to BMP signaling, a mechanism that might be required to buffer spatial fluctuations in *bmp* expression within the gastrula. Such function Nodal signaling in repressing BMP signaling is also supported by our findings in embryos, where axis elongation defects upon *bmp* overexpression are more pronounced if peak Nodal signaling levels are reduced, and in explants, where BMP and Nodal signaling form opposing gradients with Nodal signaling attenuating BMP signaling activity directly at the site of presumptive explant elongation. Why the impact of Nodal signaling becomes more apparent in the explant context than the embryo is still unclear, but likely signaling from the extraembryonic yolk cell (Sun *et al*, 2014) and/or embryo geometry contribute in maintaining the BMP signaling domain in the embryo in absence of Nodal signaling. Notably, opposing gradients of Nodal and BMP signaling activity have recently also been shown to spontaneously arise in elongating 3D mESC gastruloids (McNamara *et al*, 2023), suggesting that their coordinated spatial organization might also be involved in gastruloid morphogenesis.

It has recently been proposed that spatially unconstrained *in vitro* gastrulation models adopt a universally conserved elongated shape that might represent a ‘ground state’ of development (Steventon *et al*, 2021; Anlas & Trivedi, 2021). Consistent with this, zebrafish embryonic explants have previously been shown to elongate upon local Nodal signaling activation (Xu *et al*, 2014; Fulton *et al*, 2020; Schauer *et al*, 2020; Williams & Solnica-Krezel, 2020b; Cheng *et al*, 2023), a behavior we now show to be predominantly mediated by medially-oriented mesendodermal cell intercalations. As Nodal signaling can in principle also trigger ectodermal cell intercalations in zebrafish explants (Williams & Solnica-Krezel, 2020b), future work will have to investigate how the integration of mesendodermal and ectodermal cell dynamics promotes overall axis elongation. In contrast to the prevalence of oriented mesendoderm intercalations during explant elongation, C&E movements in the mesendoderm of intact zebrafish embryos require a combination of collective migration and intercalation movements to drive body axis elongation (Tada & Heisenberg, 2012). Indeed, increasing the population of mesendodermal progenitors associated with migratory behavior by overactivating BMP signaling diminishes body axis elongation both in explants and in the intact embryo (Myers *et al*, 2002a; von der Hardt *et al*, 2007). This supports the notion that cell intercalations rather than just migration are a prerequisite for effective body axis elongation. Considering previous work showing that mediolateral cell intercalations drive *in vitro* body axis elongation in *Xenopus* tissue explants (Huebner & Wallingford, 2018; Keller & Sutherland, 2020), our results suggest that cell intercalations constitute a conserved mechanism for elongating mesenchymal embryonic tissues independently of the specific embryonic context. In this process, BMP signaling needs to be tightly spatially controlled, likely for segregating the domain of cell migration, leading to the accumulation of cells on the dorsal side, versus dorsal cell intercalation, elongating the body axis. As ectopic BMP signaling also interferes with effective elongation of *Xenopus* explants and embryos, as well as stem cell-derived gastruloids (Graff *et al*, 1994; De Robertis *et al*, 2017; Turner *et al*, 2017; Yoon *et al*, 2021), such potential balancing function of migration and intercalation could be conserved. Further identification of molecular and cellular effectors by which BMP signaling changes the morphogenetic capacity of cells in different organisms will thus be key to generate important insights into the conserved and species-specific functions of BMP signaling to shape the gastrula stage embryo.

Properly shaping embryonic tissues requires a large and versatile morphogenetic toolkit, as embryonic morphologies can vary widely between species. For instance in zebrafish embryos, which spread over a big extraembryonic yolk cell in the course of gastrulation, mesendoderm is induced in a narrow band of cells all around the germ ring margin, whereas in zebrafish explants, lacking a yolk cell, mesendoderm arises from a local, compact domain (Feldman *et al*, 1998; Rebagliati *et al*, 1998; Fan *et al*, 2007; Harvey & Smith, 2009; Dubrulle *et al*, 2015; van Boxtel *et al*, 2015; Rogers *et al*, 2017; Schauer *et al*, 2020; Fulton *et al*, 2020; Pinheiro *et al*, 2022). This geometrical constraint potentially affects the function of BMP signaling, capable of forming a long-range activity gradient, in balancing cell migration versus intercalation. It is conceivable that the function of BMP signaling activity in suppressing premature cell intercalations becomes particularly important in embryos where mesendoderm specification is spread over long distances and thus cells need to undergo a long-range convergence movement before arriving at the dorsal side where cell intercalations are needed for body axis elongation. Future work will have to address how the activity of BMP signaling during gastrulation has been adapted to the specific embryo geometry in different species to ensure the proper balance between convergent cell migration and dorsal cell intercalation needed for body axis formation.

## Acknowledgements

We thank Patrick Müller for sharing the *chordin*^tt250^ mutant zebrafish line as well as the plasmid for *chrd-GFP*, Katherine Rogers for sharing the *bmp2b* plasmid and Andrea Pauli for sharing the *draculin* plasmid. Diana Pinheiro generated the MZ*lefty1,2*;Tg(*sebox::EGFP*) line. We are grateful to Patrick Müller, Diana Pinheiro and Katherine Rogers and members of the Heisenberg lab for discussions, technical advice and feedback on the manuscript. We also thank Anna Kicheva and Edouard Hannezo for discussions. We thank the Imaging and Optics Facility as well as the Life Science facility at IST Austria for support. This work was supported by an ERC Advanced Grant to C.-P.H. (MECSPEC 742573). A.S. is a recipient of a DOC Fellowship of the Austrian Academy of Science at IST Austria.

## Author contributions

A.S. and C.-P.H. designed the research. A.S. performed most experiments and analyzed the experimental data. K.P.-F. performed qRT-PCRs and genotyping. R.H. wrote the code to analyze the cell dispersal data. A.S. and C.-P.H. wrote the manuscript with input from all authors.

## Supplementary figure legends

**Figure S1.**
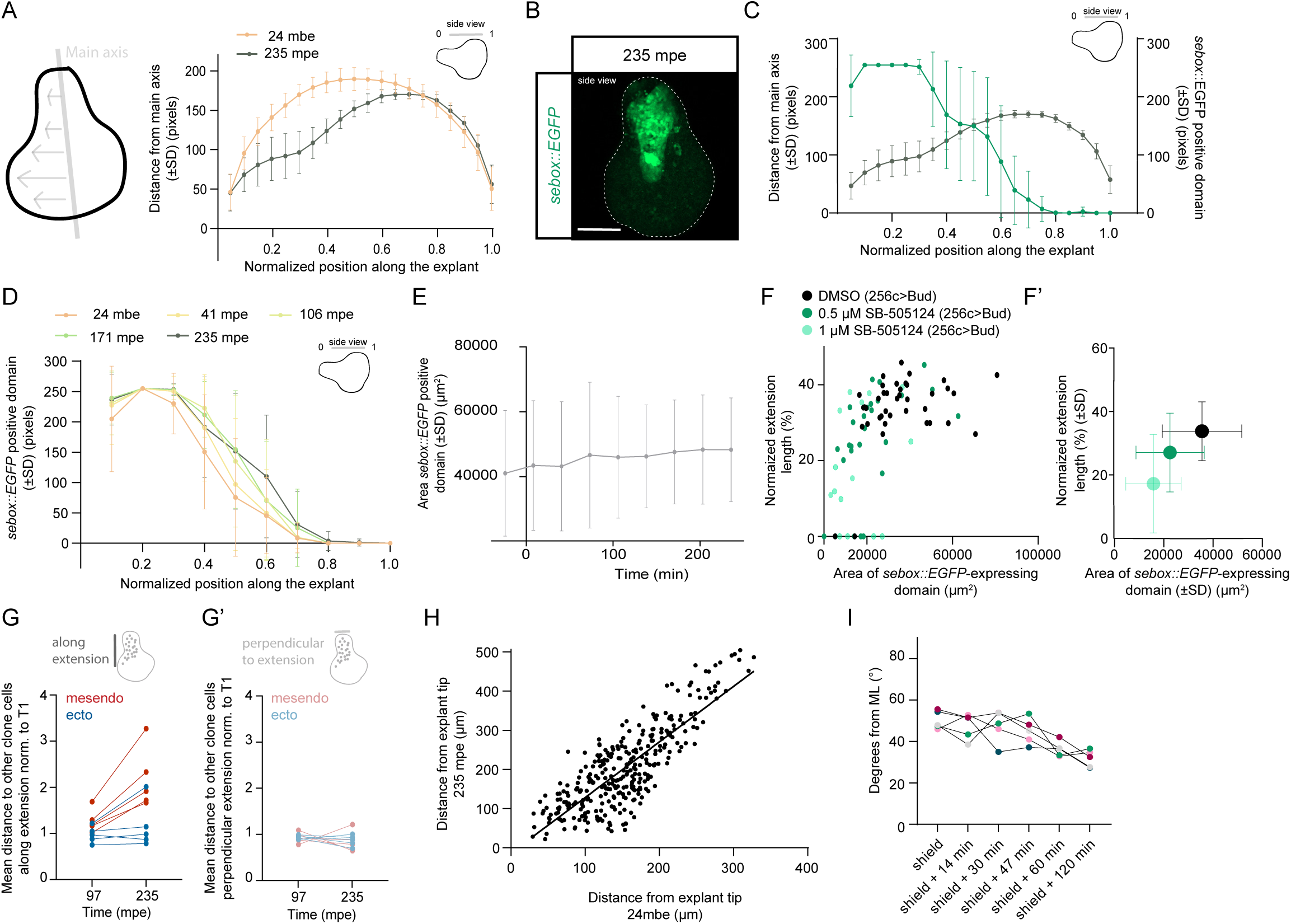
Characterization of blastoderm explant elongation. (**A**) Blastoderm explant shape assessed by the distance from the rim to the middle axis of the explant 24 min before the onset of explant elongation (mbe) and 235 minutes post explant elongation onset (mpe) (n=6, N=6). (**B**) Maximum intensity projection of fluorescence images (side views) of blastoderm explants obtained from Tg(*sebox::EGFP*) embryos expressing EGFP (green) in mesendoderm progenitors at 235 mpe. (**C**) Blastoderm explant shape assessed by the distance from the rim to the middle axis of the explant at 235 mpe and frequency of EGFP expression in blastoderm explants obtained from Tg(*sebox::EGFP*) embryos illustrating the position of mesendodermal tissues along the blastoderm explant axis (n=6, N=6). (**D**) Likelihood of EGFP expression as a function of the normalized position along the explant main axis in blastoderm explants obtained from Tg(*sebox::EGFP*) embryos, illustrating the position of mesendodermal tissues along the blastoderm explant axis at different timepoints of explant elongation (n=6, N=6). (**E**) Area of the EGFP expression domain in blastoderm explants obtained from Tg(*sebox::EGFP*) embryos at different timepoints (n=6, N=6). (**F,F’**) Area of the EGFP expression domain in blastoderm explants versus the normalized extension length in explants obtained from Tg(*sebox::EGFP*) embryos treated with DMSO (treated from 256c to bud: n=39, N=4) or Nodal inhibitor (SB-505124; treated from 256c to bud with 0.5 µM SB-505124: n=33, N=4; or 1 µM SB-505124: n=30, N=4) at bud stage. (F). (**F’**) Area of the EGFP expression domain in blastoderm explants obtained from Tg(*sebox::EGFP*) embryos versus the normalized extension length, shown as mean +/- SD, treated with DMSO (treated from 256c to bud: n=39, N=4) or Nodal inhibitor (SB-505124; treated from 256c to bud with 0.5 µM SB-505124: n=25, N=4; or 1 µM SB-505124: n=16, N=4) at bud stage. Notably, only explants which have formed mesendoderm, as assessed by Tg(*sebox::EGFP*) expression, were considered. (**G,G’**) Clone dispersal parallel (G) and perpendicular (G’) to the axis of explant elongation assessed by the mean distance of each cell in the clone to other clone cells at two timepoints during explant elongation (97 mpe and 235 mpe) for each individual explant corresponding to Fig. 1H,I (n=5, N=5). (**H**) Correlation between the position of clonally labeled cell nuclei in the mesendoderm before the onset of explant elongation (24 mbe) and during explant elongation (235 mpe) relative to the tip of the extension in wildtype explants (R^2^=0.6748, N=5). (**I**) Mean cell alignment assessed by the deviation (degrees) of the main cell extension axis from the main mediolateral explant axis at the onset of explant elongation (shield) and during explant elongation (shield + 14 min, shield + 30 min, shield + 47 min, shield + 60 min, shield + 120 min) for each individual explant corresponding to Fig. 1K (n=5, N=5). Each color indicates a different explant. Scale bars: 200 µm (B).

**Figure S2.**
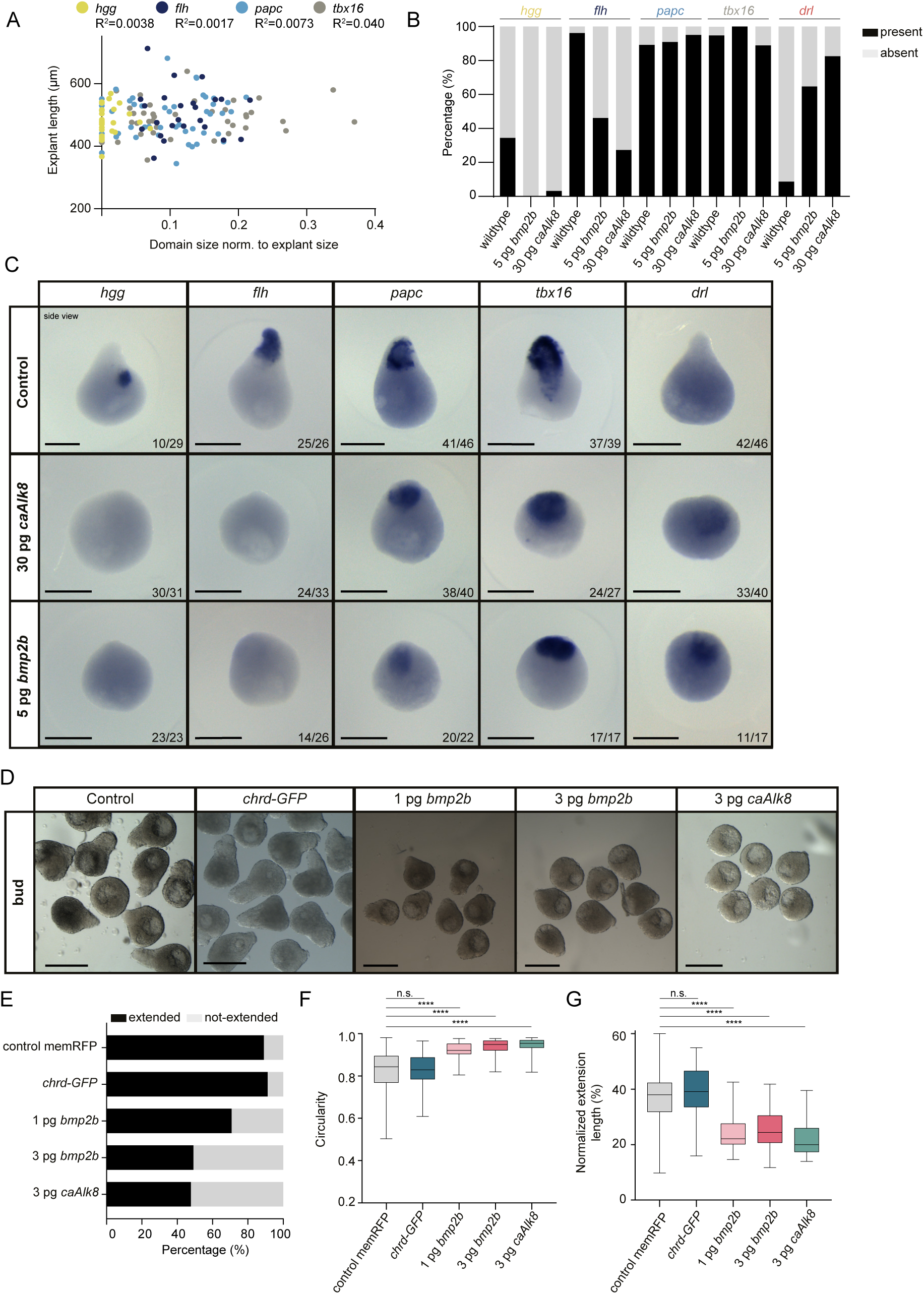
Changes in mesendoderm marker expression and blastoderm explant morphogenesis upon BMP signaling alterations. (**A**) Relationship between the length of blastoderm explants and the size of mesendodermal marker expression as assessed by *in situ* hybridization normalized to explant size at bud stage (*hgg*: n=29, N=3; *flh*: n=24, N=3; *papc*: n=47, N=5; *tbx16*: n=39, N=4). (**B**) Percentage of blastoderm explants from wildtype embryos, and embryos injected with 5pg *bmp2b* or 30pg *caAlk8* mRNA at bud stage expressing/not expressing prechordal plate (*hgg*: wildtype: n=29, N=3; 5pg *bmp2b*: n=23, N=2; 30pg *caAlk8*: n=31, N=3), notochord (*flh*: wildtype: n=26, N=3; 5pg *bmp2b*: n=26, N=2; 30pg *caAlk8*: n=33, N=3), paraxial mesoderm (*papc*: wildtype: n=46, N=5; 5pg *bmp2b*: n=22, N=2; 30pg *caAlk8*: n=40, N=4), ventrolateral mesoderm (*tbx16*: wildtype: n=39, N=4; 5pg *bmp2b*: n=17, N=2; 30pg *caAlk8*: n=27, N=4) or ventral mesoderm (*drl*: wildtype: n=46, N=3; 5pg *bmp2b*: n=17, N=2; 30pg *caAlk8*: n=40, N=3) markers assessed by *in situ* hybridization. All embryos were co-injected with 50-100pg *memRFP* as injection control. (**C**) Expression of prechordal plate (*hgg*), notochord (*flh*), paraxial mesoderm (*papc*), ventrolateral mesoderm (*tbx16*) and ventral mesoderm (*drl*) markers assessed by *in situ* hybridization in blastoderm explants from wildtype embryos, and embryos injected with 5pg *bmp2b* or 30pg *caAlk8* mRNA (side views) at bud stage. Images correspond to quantifications in (B). The proportion of explants with a similar expression pattern to the shown image is indicated in the lower right corner. (**D**) Single-plane bright-field images of blastoderm explants from wildtype embryos (control: n=236, N=13), and embryos injected with 37pg *chrd-GFP* (n=45, N=4), 1pg *bmp2b* (n=41, N=5), 3pg *bmp2b* (n=59, N=5) or 3pg *caAlk8* mRNA (n=44, N=4) (side views) at bud stage. Control explants partially correspond to explants shown in Fig. 2A. All embryos were co-injected with 50-100pg *memRFP* or *memGFP* as injection control. (**E**) Percentage of extended/not-extended explants from wildtype embryos (control: n=236, N=13), and embryos injected with 37pg *chrd-GFP* (n=45, N=4), 1pg *bmp2b* (n=41, N=5), 3pg *bmp2b* (n=59, N=5) or 3pg *caAlk8* mRNA (n=44, N=4) at bud stage. Control explants partially correspond to explants shown in Fig. 2B. All embryos were co-injected with 50-100pg *memRFP* or *memGFP* as injection control. (**F**) Circularity of extended/not-extended blastoderm explants from wildtype embryos (control: n=236, N=13), and embryos injected with 37pg *chrd-GFP* (n=45, N=4), 1pg *bmp2b* (n=41, N=5), 3pg *bmp2b* (n=59, N=5) or 3pg *caAlk8* mRNA (n=44, N=4) at bud stage. Control explants partially correspond to explants shown in Fig. 2C. All embryos were co-injected with 50-100pg *memRFP* or *memGFP* as injection control. ****p<0.0001, ns, not significant (Kruskal-Wallis test). (**G**) Normalized extension length of extended blastoderm explants from wildtype embryos (control: n=210, N=13), and embryos injected with 37pg *chrd-GFP* (n=41, N=4), 1pg *bmp2b* (n=29, N=5), 3pg *bmp2b* (n=29, N=5) or 3pg *caAlk8* mRNA (n=21, N=4) at bud stage. Control explants partially correspond to explants shown in Fig. 2D. All embryos were co-injected with 50-100pg *memRFP* or *memGFP* as injection control. ****p<0.0001, ns, not significant (Kruskal-Wallis test). Scale bars: 250 µm (C), 500 µm (D).

**Figure S3.**
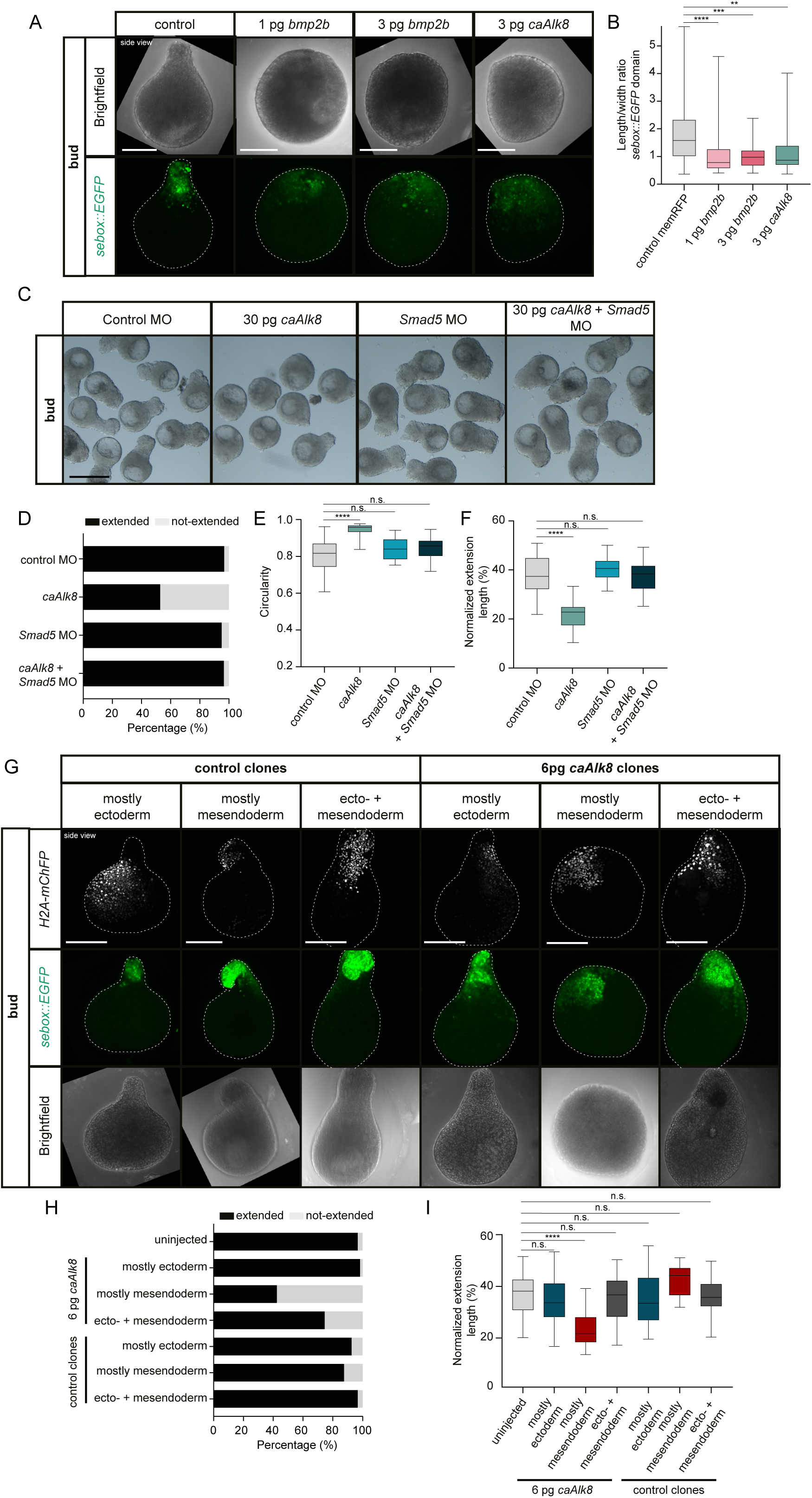
Changes in blastoderm explant length upon BMP signaling overactivation. (**A**) Maximum intensity projection of fluorescence images (side views) of blastoderm explants from Tg(*sebox::EGFP*) wildtype embryos expressing EGFP (green) in mesendoderm progenitors (control: n=62, N=8) and Tg(*sebox::EGFP*) embryos injected with 1pg *bmp2b* (n=31, N=6), 3pg *bmp2b* (n=25, N=6) and 3pg *caAlk8* (n=21, N=4) at bud stage. Control explants correspond to explants shown in Fig. 2E. All embryos were co-injected with 50-100pg *memRFP* as injection control. (**B**) Length/width ratio of the EGFP expression domain in blastoderm explants from Tg(*sebox::EGFP*) wildtype embryos marking mesendoderm progenitors (control: n=62, N=8) and Tg(*sebox::EGFP*) embryos injected with 1pg *bmp2b* (n=31, N=6), 3pg *bmp2b* (n=25, N=6) and 3pg *caAlk8* (n=21, N=4) at bud stage. Control explants correspond to explants shown in Fig. 2F. All embryos were co-injected with 50-100pg *memRFP* as injection control. ****p<0.0001, ***p=0.0008, **p=0.0032 (Kruskal-Wallis test). (**C**) Single-plane bright-field images of blastoderm explants from wildtype embryos (controlMO: n=31, N=3) and embryos injected with 30pg *caAlk8* (n=36, N=3), *Smad5*MO (n=20, N=3) and 30pg *caAlk8* + *Smad5*MO (n=29, N=3) (side views) at bud stage. All embryos were co-injected with 50-100pg *memRFP* or *memGFP* as injection control. (**D**) Percentage of extended/not-extended blastoderm explants from wildtype embryos (controlMO: n=31, N=3) and embryos injected with 30pg *caAlk8* (n=36, N=3), *Smad5*MO (n=20, N=3) and 30pg *caAlk8* + *Smad5*MO (n=29, N=3) at bud stage. All embryos were co-injected with 50-100pg *memRFP* or *memGFP* as injection control. (**E**) Circularity of extended or not-extended blastoderm explants from wildtype embryos (controlMO: n=31, N=3) and embryos injected with 30pg *caAlk8* (n=36, N=3), *Smad5*MO (n=20, N=3) and 30pg *caAlk8* + *Smad5*MO (n=29, N=3) at bud stage. All embryos were co-injected with 50-100pg *memRFP* or *memGFP* as injection control. ****p<0.0001, ns, not significant (Kruskal-Wallis test). (**F**) Normalized extension length of extended blastoderm explants from wildtype embryos (controlMO: n=30, N=3) and embryos injected with 30pg *caAlk8* (n=18, N=3), *Smad5*MO (n=19, N=3) and 30pg *caAlk8* + *Smad5*MO (n=29, N=3) at bud stage. All embryos were co-injected with 50-100pg *memRFP* or *memGFP* as injection control. ****p<0.0001, ns, not significant (One-way ANOVA). (**G**) Maximum intensity projection of fluorescence images (side views) of blastoderm explants obtained from Tg(*sebox::EGFP*) embryos expressing EGFP (green) in mesendoderm progenitors at bud stage showing clonally labeled cell nuclei of control blastomere injections (5pg *H2A-mChFP* mRNA injected) or blastomere injections with 6pg *caAlk8* and 5pg *H2A-mChFP* mRNA (grey) positioned either mostly in the ectoderm (control: n=27, N=5; 6pg *caAlk8*: n=57, N=5), mostly in the mesendoderm (control: n=8, N=5; 6pg *caAlk8*: n=33, N=5) or in both the ectoderm and mesendoderm (control: n=30, N=5; 6pg *caAlk8*: n=47, N=5). (**H**) Percentage of extended/not-extended blastoderm explants obtained from uninjected wildtype embryos (n=60, N=5), single blastomere injected embryos with 5pg *H2A-mChFP* (positioned mostly in the ectoderm: n=27, N=5; mostly in the mesendoderm: n=8, N=5; in both the ectoderm and mesendoderm: n=30, N=5) and single blastomere injected embryos with 5pg *H2A-mChFP* + 6pg *caAlk8* positioned mostly in the ectoderm: n=57, N=5; mostly in the mesendoderm: n=33, N=5; in both the ectoderm and mesendoderm: n=47, N=5) blastoderm explants at bud stage. (**I**) Normalized extension length of extended blastoderm explants obtained from uninjected wildtype embryos (n=58, N=5), single blastomere injected embryos with 5pg *H2A-mChFP* (positioned mostly in the ectoderm: n=25, N=5; mostly in the mesendoderm: n=7, N=5; in both the ectoderm and mesendoderm: n=29, N=5) and single blastomere injected embryos with 5pg *H2A-mChFP* + 6pg *caAlk8* (mostly in the ectoderm: n=56, N=5; mostly in the mesendoderm: n=14, N=5; in both ectoderm and mesendoderm: n=35, N=5) at bud stage. ****p<0.0001, ns, not significant (One-way ANOVA). Scale bars: 200 µm (A,G), 500 µm (C).

**Figure S4.**
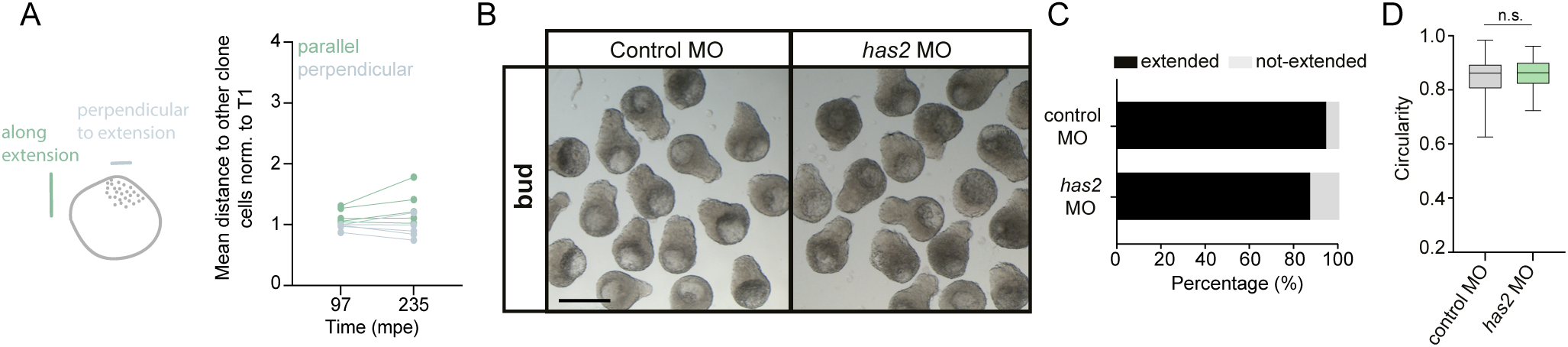
Elongation of blastoderm explants upon *has2* knock-down. (**A**) Clone dispersal parallel (dark green) and perpendicular (light green) to the axis of explant elongation assessed by the mean distance of each cell in the clone to other clone cells at two timepoints during explant elongation (97 mpe and 235 mpe) for each individual 30pg *caAlk8* explant. (**B**) Single-plane bright-field images (side views) of blastoderm explants from wildtype embryos (controlMO: n=102, N=6) and embryos injected with *has2*MO (n=91, N=6) at bud stage. All embryos were co-injected with 50-100pg *memRFP* or *memGFP* as injection control. (**C**) Percentage of extended/not-extended blastoderm explants from wildtype embryos (controlMO: n=102, N=6) and embryos injected with *has2*MO (n=91, N=6) at bud stage. All embryos were co-injected with 50-100pg *memRFP* or *memGFP* as injection control. (**D**) Circularity of extended or not-extended blastoderm explants from wildtype embryos (controlMO n=102, N=6) and embryos injected with *has2*MO (n=91, N=6) at bud stage. All embryos were co-injected with 50-100pg *memRFP* or *memGFP* as injection control. ns, not significant (Unpaired t-test). Scale bar: 500 µm (B).

**Figure S5.**
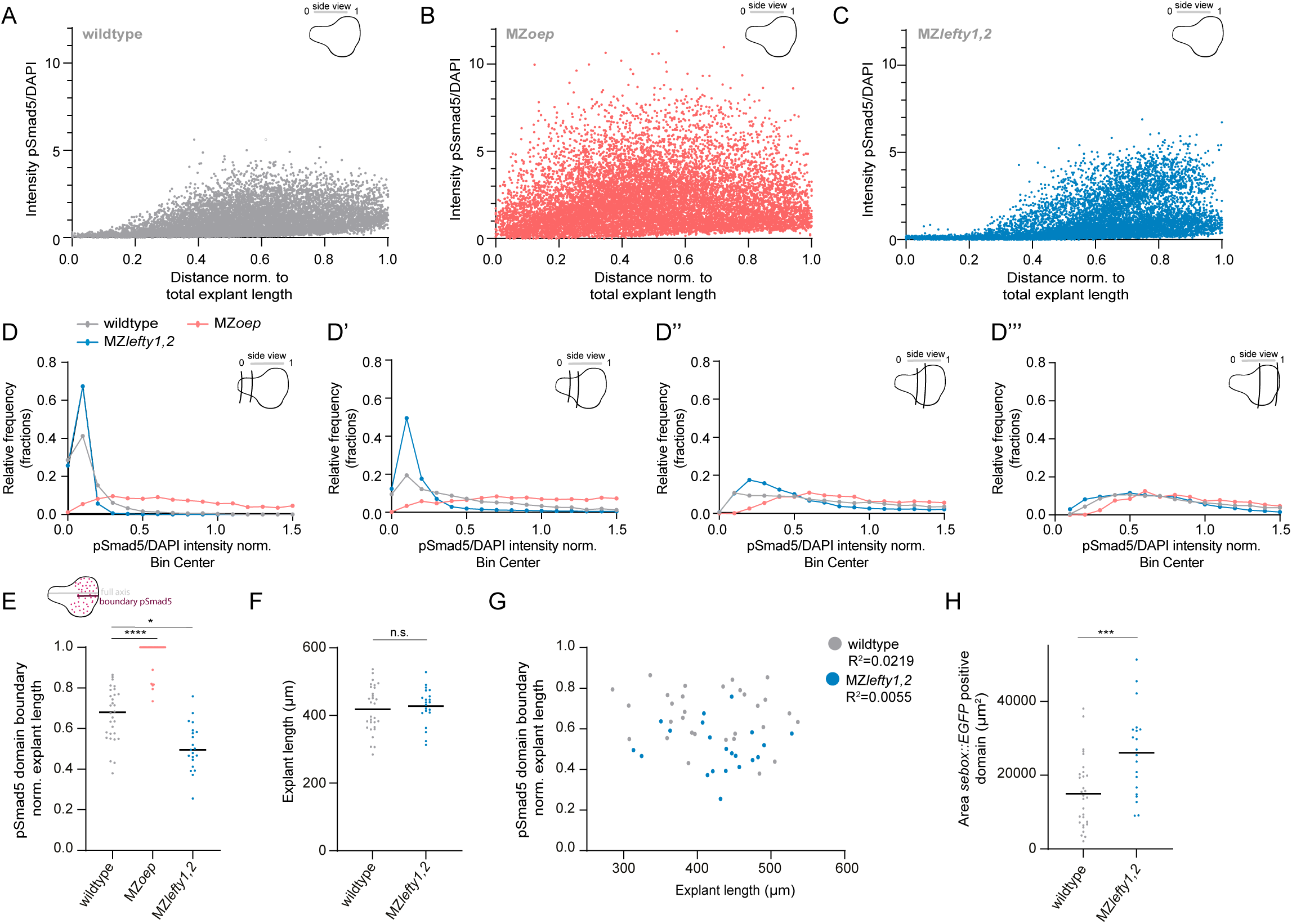
Profile of pSmad5 distribution upon perturbation of Nodal signaling. (**A**) Intensity of nuclear pSmad5 normalized to DAPI as a function of the distance from the explant tip in blastoderm explants from wildtype embryos (n=9, N=4) during explant elongation (corresponding to embryonic 75% epiboly stage). (**B**) Intensity of nuclear pSmad5 normalized to DAPI as a function of the distance from the explant tip in blastoderm explants from MZ*oep* embryos (n=9, N=4) during explant elongation (corresponding to embryonic 75% epiboly stage). (**C**) Intensity of nuclear pSmad5 normalized to DAPI as a function of the distance from the explant tip in blastoderm explants from MZlefty*1,2* embryos (n=9, N=4) during explant elongation (corresponding to embryonic 75% epiboly stage). (**D-D’’’**) Frequency distribution of pSmad5/DAPI ratio relative to the mean intensity in the bin closest to the back (high intensity domain) of the wildtype explants, as determined in Fig. 4F, in discrete spatial bins across the tip-back axis for blastoderm explants from wildtype (grey), MZ*oep* (salmon) and MZ*lefty1,2* (blue) mutant embryos during explant elongation (corresponding to embryonic 75% epiboly stage). Notably the first and last 4% of the explant were excluded due to low number of nuclei at the ends. The spatial bins are thus shown as (D) 0.04-0.25 from the tip, (D’) 0.25-0.5 from the tip, (D’’) 0.5-0.75 from the tip, (D’’’) 0.75-0.96 from the tip. (**E**) Domain extent of nuclear pSmad5 measured centrally from the back of the explant normalized to the explant length in blastoderm explants obtained from wildtype embryos (n=30, N=5), MZ*oep* embryos (n=31, N=5) and MZ*lefty1,2* embryos (n=21, N=5) during explant elongation (corresponding to embryonic 75% epiboly stage) with the black line indicating the median. ****p<0.0001, *p=0.028 (Kruskal-Wallis test). (**F**) Length of blastoderm explants obtained from Tg(*sebox::EGFP*) embryos (n=30, N=5) and MZ*lefty1,2*;Tg(*sebox::EGFP*) embryos (n=21, N=5) during explant elongation (corresponding to embryonic 75% epiboly stage) with the black line indicating the mean. ns, not significant (Unpaired t-test). (**G**) Relationship between the length of blastoderm explants obtained from Tg(*sebox::EGFP*) embryos (n=30, N=5) and MZ*lefty1,2*; Tg(*sebox::EGFP*) embryos (n=21, N=5) during explant elongation (corresponding to embryonic 75% epiboly stage) and the pSmad5 domain extent normalized to the whole explant length. (**H**) Area of the EGFP expression domain marking mesendodermal progenitors normalized to the overall explant area in blastoderm explants obtained from Tg(*sebox::EGFP*) embryos (n=30, N=5) and MZ*lefty1,2*;Tg(*sebox::EGFP*) embryos (n=20, N=5) during explant elongation (corresponding to embryonic 75% epiboly stage) with the black line indicating the mean. ***p=0.0006 (Unpaired t-test).

**Figure S6.**
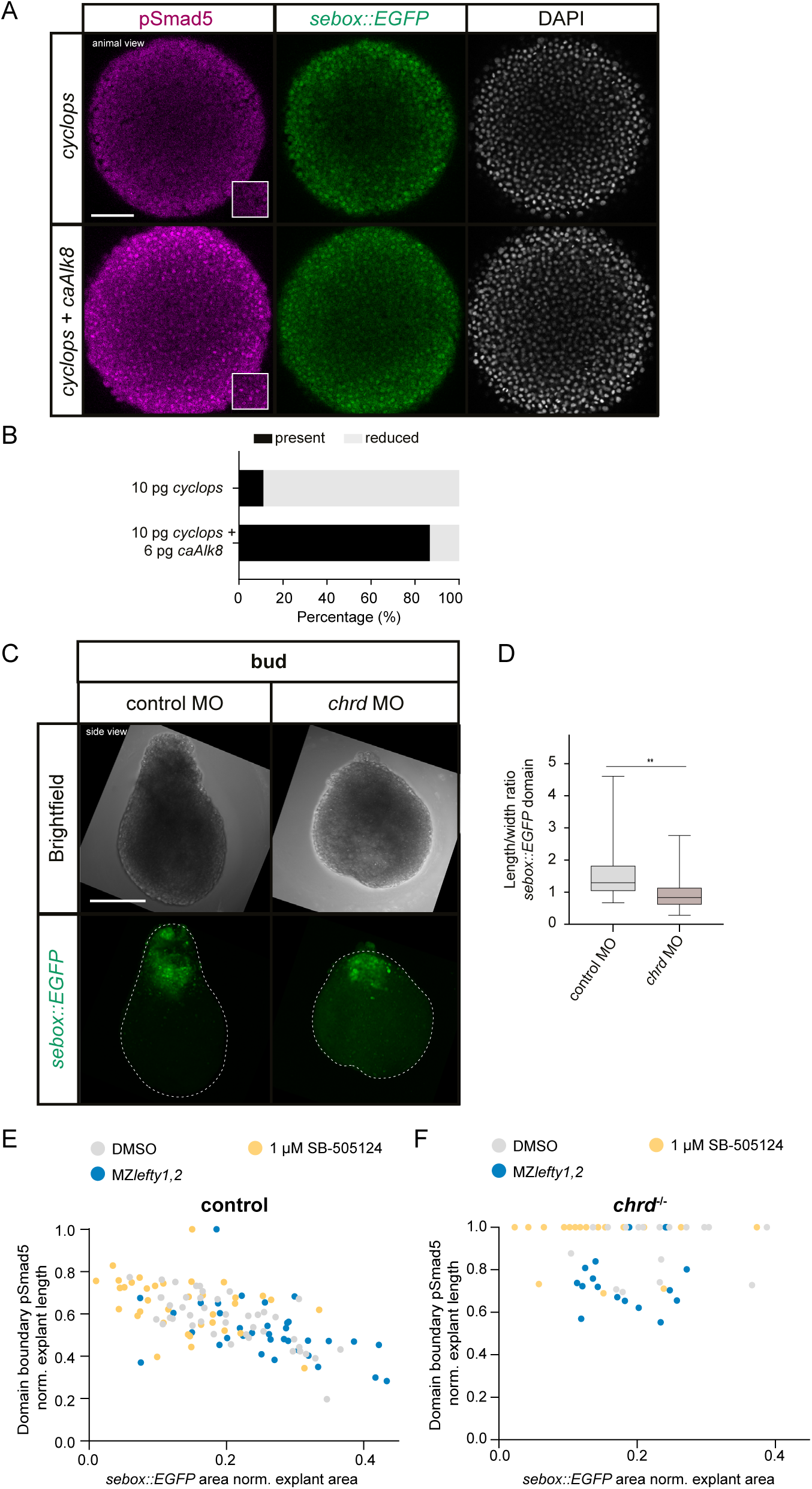
Repression of BMP signaling activity by Nodal-dependent regulation of *chordin* expression. (**A**) Single-plane high-resolution images (animal view) of Tg(*sebox::EGFP*) embryos marking mesendoderm progenitors (green) at 75% epiboly stained for pSmad5 (BMP signaling activity; magenta) and DAPI (nuclei; grey) injected with 10pg *cyclops* (top) (n=18, N=3) or 10pg *cyclops* + 6pg *caAlk8* mRNA (bottom) (n=15, N=3). (**B**) Percentage of present/absent nuclear pSmad5 in embryos injected with 10pg *cyclops* (n=18, N=3) or 10pg *cyclops* + 6pg *caAlk8* mRNA (n=15, N=3) at 75% epiboly stage. (**C**) Maximum intensity projection of fluorescence images (side views) of blastoderm explants from Tg(*sebox::EGFP*) embryos expressing EGFP (green) in mesendoderm progenitors (5ng controlMO: n=26, N=3) and embryos injected with 4.5ng *chrd*MO (n=17, N=3) at bud stage. All embryos were co-injected with 50-100pg *memRFP* as injection control. (**D**) Length/width ratio of the EGFP expression domain in blastoderm explants from Tg(*sebox::EGFP*) wildtype embryos marking mesendoderm progenitors (5ng controlMO: n=26, N=3) and embryos injected with 4.5ng *chrd*MO (n=17, N=3) at bud stage. All embryos were co-injected with 50-100pg *memRFP* as injection control. **p=0.0013 (Mann-Whitney test). (**E**) Domain of nuclear pSmad5 measured centrally from the back of the explant, normalized to the explant length, and area of *sebox::EGFP* expression, normalized to explant area, in blastoderm explants obtained from wildtype embryos treated with DMSO (treated from 256c to 75% epiboly: wildtype: n=43, N=6) or 1 µM Nodal inhibitor (SB-505124; treated from 256c to 75% epiboly: wildtype: n=36, N=6) and MZ*lefty1,2* (n=35, N=5) embryos during explant elongation (corresponding to embryonic 75% epiboly stage) with Tg(*sebox::EGFP*) marking mesendoderm progenitors. (**F**) Domain of nuclear pSmad5 measured centrally from the back of the explant, normalized to the explant length, and area of *sebox::EGFP* expression, normalized to explant area, in blastoderm explants obtained from *chrd*^-/-^ embryos treated with DMSO (treated from 256c to 75% epiboly: *chrd*^-/-^: n=17, N=4) or 1 µM Nodal inhibitor (SB-505124; treated from 256c to 75% epiboly: *chrd*^-/-^: n=17, N=6) and MZ*lefty1,2;chrd*^-/-^ (n=17, N=4) embryos during explant elongation (corresponding to embryonic 75% epiboly stage) with Tg(*sebox::EGFP*) marking mesendoderm progenitors. Scale bars: 100 µm (A), 200 µm (C).

**Figure S7.**
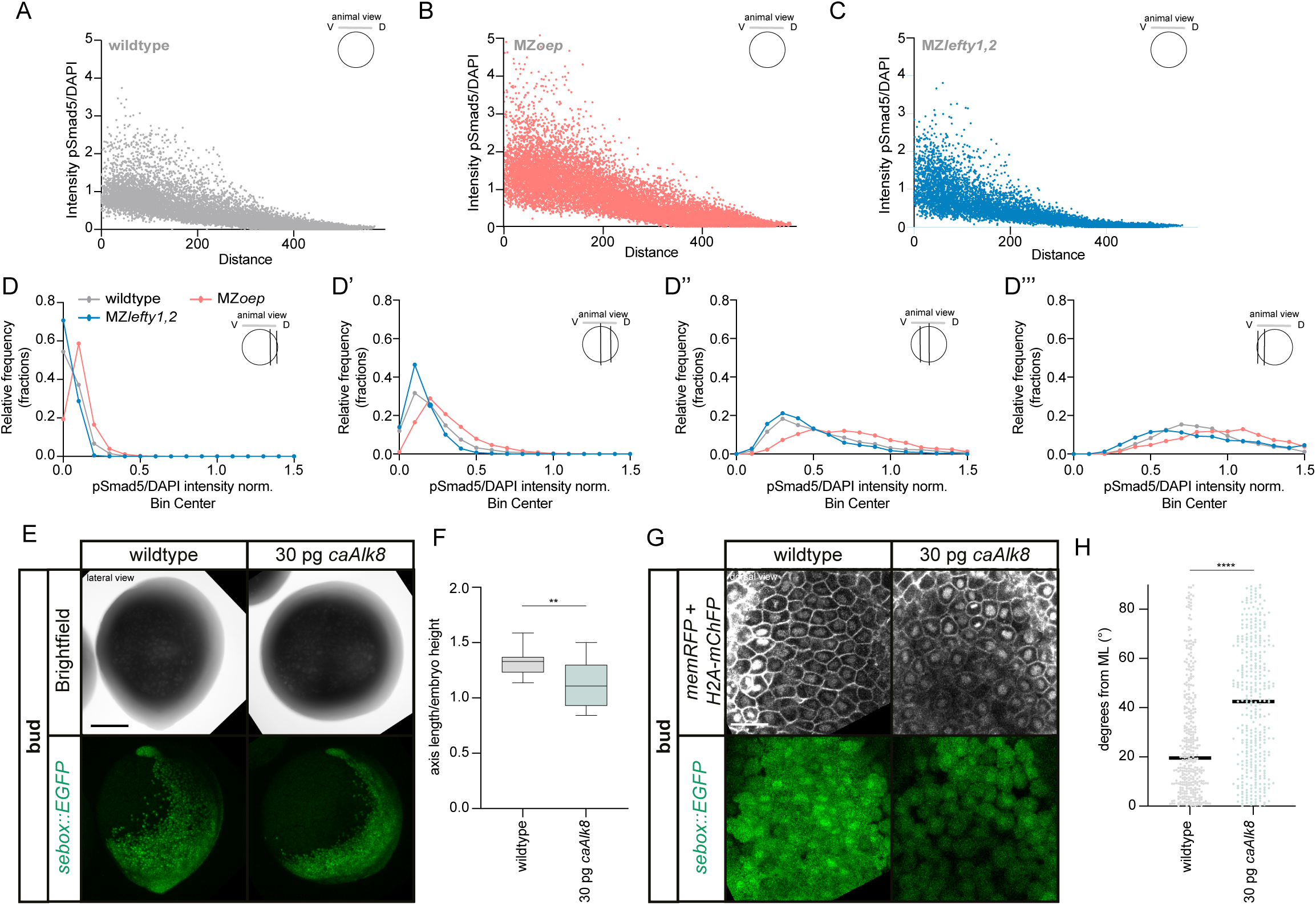
Changes in axis elongation upon BMP signaling overactivation. (**A**) Intensity of nuclear pSmad5 normalized to DAPI as a function of the distance from the ventral side in wildtype embryos (n=11, N=5) at 75% epiboly stage. D=dorsal, V=ventral. (**B**) Intensity of nuclear pSmad5 normalized to DAPI as a function of the distance from the ventral side in MZ*oep* embryos (n=11, N=5) at 75% epiboly stage. (**C**) Intensity of nuclear pSmad5 normalized to DAPI as a function of the distance from the ventral side in MZ*lefty1,2* embryos (n=11, N=5) at 75% epiboly stage. (**D-D’’’**) Frequency distribution of pSmad5/DAPI ratio relative to the mean intensity in the bin closest to the ventral side (high intensity domain) of the wildtype embryos, as determined in Fig. 6B, in discrete spatial bins across the dorsoventral axis for wildtype (grey), MZ*oep* (salmon) and MZ*lefty1,2* (blue) mutant embryos respectively. Notably the first and last 4% of the embryo length were excluded due to low number of nuclei at the sample edges. The spatial bins are thus shown as (D) 0.04-0.25 from dorsal, (D’) 0.25-0.5 from dorsal, (D’’) 0.5-0.75 from dorsal, (D’’’) 0.75-0.96 from dorsal. (**E**) Maximum intensity projection of bright-field (top) and fluorescence (bottom) images (lateral views; dorsal pointing to the right) of Tg(*sebox::EGFP*) wildtype embryos (n=16, N=5) marking mesendodermal progenitors (left) or Tg(*sebox::EGFP*) embryos injected with 30pg *caAlk8* (right) (n=17, N=5) at bud stage. All embryos were co-injected with 80pg *memRFP* or 40pg *H2A- mChFP* as injection control. (**F**) Length of the mesendodermal embryonic axis on the dorsal side over embryo height at bud stage in wildtype embryos (n=16, N=5) and embryos injected with 30pg *caAlk8* mRNA (n=17, N=5). All embryos were co-injected with 80pg *memRFP* or 40pg *H2A-mChFP* as injection control. **=0.0015 (Unpaired t-test). (**G**) Single-plane high-resolution images of wildtype and 30pg *caAlk8* injected embryos (dorsal views) expressing Tg(*sebox::EGFP*) marking mesendodermal progenitors at 90% epiboly stage. Cell outlines are marked by 80-100pg *memRFP* mRNA injection (grey) expression. Cell nuclei are marked by 40pg *H2A-mChFP* mRNA injection (grey) expression. (**H**) Cell alignment assessed by the deviation (degrees) of the main cell extension axis from the main mediolateral (ML) embryo axis during axis elongation at 90% epiboly (wildtype: 362 cells, n=7, N=4; 30pg *caAlk8*: 357 cells, n=7, N=4). ****p<0.0001 (Mann-Whitney test). Scale bars: 200 µm (E), 100 µm (G).

## Supplementary Video legends

**Supplementary Video 1. Blastoderm and mesendoderm morphogenesis in wildtype explants.** Maximum intensity projection of fluorescence/brightfield high-resolution time-lapse imaging of blastoderm explants obtained from Tg(*sebox::EGFP*) embryos expressing EGFP (green) in mesendoderm progenitors from 24 minutes before the onset of extension (mbe). Time is shown in minutes. Scale bar: 100 µm.

**Supplementary Video 2. Mesendodermal clone dispersal in wildtype explants.** Maximum intensity projection of fluorescence/brightfield high-resolution time-lapse imaging of blastoderm explants showing tracked clonally labeled cell nuclei in the mesendoderm (red) from 24 minutes before the onset of extension (mbe). Time is shown in minutes. Scale bar: 100 µm.

**Supplementary Video 3. Mesendodermal cell alignment in wildtype explants.** Maximum intensity projection of medium planes of high-resolution images of blastoderm explants (side views) obtained from Tg(*sebox::EGFP*) embryos expressing EGFP in mesendoderm progenitors at the onset of explant elongation (corresponding to embryonic shield stage). Mesendodermal cell boundaries within the explant extension are marked by clonal lifeact-RFP (grey) expression. Time is shown in minutes. Scale bar: 100 µm.

**Supplementary Video 4. Ectodermal clone dispersal in wildtype explants.** Maximum intensity projection of fluorescence/brightfield high-resolution time-lapse imaging of blastoderm explants showing tracked clonally labeled cell nuclei in the ectoderm (blue) from 24 minutes before the onset of extension (mbe). Time is shown in minutes. Scale bar: 100 µm.

**Supplementary Video 5. Mesendodermal clone dispersal in *caAlk8* overexpressing explants.** Maximum intensity projection of fluorescence/brightfield high-resolution time-lapse imaging of blastoderm explants showing tracked clonally labeled cell nuclei in the mesendoderm (red) of explants prepared from embryos overexpressing 30pg *caAlk8* from 24 minutes before the onset of extension (mbe). Time is shown in minutes. Scale bar: 100 µm.

## Supplementary Files

**Supplementary File 1.** MATLAB script for cell dispersal analysis.

## Materials and methods

### Fish lines and husbandry

Maintenance and handling of zebrafish (*Danio rerio*) was performed as previously described (Westerfield, 2000). The following zebrafish strains were used in this study: wild type AB or a cross of wild type ABxTL, Tg(*sebox*::*EGFP*) (Ruprecht *et al*, 2015), MZ*oep* (Gritsman *et al*, 1999), MZ*lefty1,2* (Rogers *et al*, 2017), MZ*oep*;Tg(*gsc::EGFP-CAAX*), MZ*lefty1,2*;Tg(*sebox*::*EGFP*), *chordin*^tt250^ (Hammerschmidt *et al*, 1996), *chordin*^tt250^; Tg(*sebox*::*EGFP*), MZ*lefty1,2*;*chordin*^tt250^;Tg(*sebox*::*EGFP*) and MZ*lefty1,2*; *chordin*^tt250^. Mutant transgenic lines were generated by crosses of the Tg(*sebox*::*EGFP*) and Tg(*gsc::EGFP-CAAX*) line with the respective mutants. Raising and genotyping of MZ*oep,* MZ*lefty1,2* and *chordin*^tt250^ was performed as described (Gritsman *et al*, 1999; Rogers *et al*, 2017; Pomreinke *et al*, 2017). Embryos were raised in E3 medium or Danieau’s solution (58 mM NaCl, 0.7 mM KCl, 0.4 mM MgSO_4_, 0.6 mM Ca(NO_3_)_2_, 5 mM HEPES, pH 7.6) at 25-31°C and staged according to (Kimmel *et al*, 1995).

### Blastoderm explant preparation

Blastoderm explants were prepared as previously described (Schauer *et al*, 2020). In short, the entire blastoderm was removed from the yolk cell at 256-cell stage using forceps and cultured at 25-31°C in Danieau’s solution. Staging of the explants was performed based on sibling embryos from the same egg lay. After approximately 1 hour, explants which did not close up properly or showed delayed cleavages, were removed. A stereo-microscope (Olympus SZX 12) with a QImaging Micropublisher 5.0 camera was used to take bright-field images of explants for analyzing explant morphology.

### Embryo microinjections

To synthesize mRNAs, the mMessage mMachine Kit (Ambion) was used. 1-cell stage and 32-128-cell stage injections were performed as described (Westerfield, 2000). The following mRNAs were used: 30-50 pg *H2A-mChFP* (Arboleda-Estudillo *et al*, 2010), 50-100 pg *membrane-RFP* (Iioka *et al*, 2004), 50-100 pg *membrane-GFP* (Kimmel & Meyer, 2010), 3-30 pg *constitutively active Alk8* (*caAlk8*) (Bauer *et al*, 2001), 1-5 pg *bmp2b* (Kishimoto *et al*, 1997), 37 pg *chrd-GFP* (Pomreinke *et al*, 2017) and 10 pg *cyclops* (Rebagliati *et al*, 1998). For clonally labeling nuclei and/or F-actin, 5 pg *H2A-mChFP* and/or 10 pg *LifeAct-RFP* (Behrndt *et al*, 2012) were injected into a single blastomere at the 32-128-cell stage. For overexpressing *caAlk8* in one or two blastomeres, 6 pg *caAlk8* were injected at 8-16-cell stage together with 5 pg *H2A-mChFP*. The following morpholinos were used in this study: 1.5 nl 0.25 mM *has2*MO (5’-AGCAGCTCTTTGGAGATGTCCCGTT-3’) (Bakkers *et al*, 2004; von der Hardt *et al*, 2007), 1.5 nl 0.5 mM *Smad5*MO (5’-AACAGACTAGACATGGAGGTCATAG-3’) (Lele *et al*, 2001a; von der Hardt *et al*, 2007), 1.5 ng *bmp2b*MO (5’- CGCGGACCACGGCGACCATGATC-3’) (Lele *et al*, 2001a), 4.5 ng *chordin*MO (5’-ATCCACAGCAGCCCCTCCATCATCC-3’) (Nasevicius & Ekker, 2000), and 5 ng controlMO (human β-globin MO 5’-CCTCTTACCTCAGTTACAATTTATA–3’, Gene Tools).

### Live imaging of blastoderm explants and embryos

Sample preparation for live imaging experiments at upright confocal microscopes was performed as described previously (Schauer *et al*, 2020) by mounting explants in 800 x 800 µm agarose molds (Microtissues) in Danieau’s solution. Unless indicated differently, live imaging at upright microscopes was started when control embryos had reached 50% epiboly stage and explants were oriented in side-view. Imaging was performed using a Zeiss LSM 900 upright microscope equipped with a Zeiss Plan-Apochromat 20x/1.0 water immersion objective. For high-magnification analysis of cell alignment in blastoderm explants at inverted confocal microscopes, small pieces of 800 x 800 µm agarose molds were cut and immobilized on glass-bottom dishes (35 mm, MatTek Corporation, cat.no.: P35G-1.5-14-C) with agarose. For analysis of cell alignment in embryos, the embryos were mounted in 0.7% low melting point agarose in Danieau’s solution on the same type of glass-bottom dishes with the dorsal side, as identified by *sebox:EGFP* signal, facing the objective. Imaging was started at shield stage, unless indicated differently, and performed on a Zeiss LSM880 inverted microscope with a Zeiss Plan-Apochromat 40x/1.2 water-immersion objective. In all cases, the temperature during image acquisition was set to 28.5°C.

### Whole mount *in situ* hybridization (WMISH)

Embryos for WMISHs were fixed overnight at 4°C in 4% PFA, washed 5x in PBS, transferred to 100% methanol and stored at −20°C until further processing. WMISHs with DIG-labeled antisense RNA probes were then performed as described previously (Thisse & Thisse, 2008). Antisense RNA probes were generated using Roche digoxigenin (DIG)- modified nucleotides with SP6, T7 or T3 RNA polymerase from mMessage mMachine kits (ThermoFisher, AM1344). The following RNA probes were used: *hgg* (Thisse *et al*, 1994), *flh* (Talbot *et al*, 1995), *papc* (Yamamoto *et al*, 1998), *tbx16* (Griffin *et al*, 1998) and *draculin* (Herbomel *et al*, 1999). Imaging of WMISHs was performed with a QImaging Micropublisher 5.0 camera on an Olympus SZX 12 stereo-microscope.

### Assessment of mesendoderm domain morphology in conjunction with explant morphology

To analyze mesendoderm induction and explant elongation upon reduction in Nodal signaling, explants were treated with either 0.1% DMSO (controls) or 0.5 µM or 1 µM SB-505124 (Byfield *et al*, 2004) from 256-cell stage until fixation at bud stage. To analyze BMP-dependent changes in mesendoderm morphology, BMP signaling was overactivated by injection of *caAlk8*, *bmp2b* mRNA or *chrd*MO together with *membraneRFP* or *H2A-mChFP* mRNA (as described above in ‘Embryo microinjections’) in a *sebox*::*EGFP* transgenic background. Amounts of mRNA and morpholino injected for specific experiments are indicated in the respective figure legends. Explants were raised until bud stage, then fixed in 4% PFA overnight at 4°C. To analyze explant elongation and clone localization upon local injection of 6 pg *caAlk8* and/or 5 pg *H2A-mChFP* mRNA, embryos were injected (as described above in ‘Embryo microinjections’) in 1-2 blastomeres at the 8-16-cell stage to create large clones of different localization and fixed at bud stage. The explants were then washed 5x in PBS + 0.1% Tween and mounted in 0.7% low melting point agarose in 2% agarose molds in side-view for imaging on a Zeiss LSM 800 Upright microscope equipped with a Zeiss Plan-Apochromat 20x/1.0 water immersion objective.

### Whole mount immunofluorescence (WMIF)

Immunostainings were performed similarly for blastoderm explants and intact embryos. α-pSmad5 (1:100, cat.no. 9516S, Cell Signaling Technology) whole mount immunofluorescence was performed as described (Pomreinke *et al*, 2017; Schauer *et al*, 2020). As secondary antibody, goat α-rabbit-Alexa Fluor 546 was used (1:500, cat.no. A11010, Thermo Fisher Scientific). α-pSmad5 and α-pSmad2 (1:1000, cat.no. 8828, Cell Signaling Technology) double WMIF was performed as described in (Soh *et al*, 2020) with goat α-rabbit-horseradish peroxidase (1:500, AB_2307391, Jackson ImmunoResearch) and TSA fluorescein detection of pSmad2 activity (Soh *et al*, 2020). The immunostained blastoderm explants and embryos were mounted in 2% agarose molds at a Zeiss LSM 880 upright microscope equipped with a Zeiss Plan-Apochromat 20x/1.0 water immersion objective. Explants were oriented in side-view using either the *sebox::EGFP* (for wildtype and MZ*lefty1,2*) or *gsc::EGFP-CAAX* (for MZ*oep*) signal as a marker. Embryos were oriented to be viewed from the animal pole. Imaging conditions between different samples and replicates were kept similar.

For staining of pSmad5 in absence of *chrd*, explants were prepared from either MZ*lefty1,2; chrd^tt250^* females crossed with MZ*lefty1,2;chrd^tt250^;*Tg(*sebox::EGFP*) mutant males or from crosses of *chrd*^tt250^ females and *chrd^tt250^*;Tg(*sebox::EGFP*) males treated with DMSO (control) or 1 µM SB505124 and fixed at 75% epiboly. Immunostaining was performed as described above, also adding chicken α-GFP (Soh *et al*, 2020) antibody (1:200, cat.no. GFP-1020, Aves Labs) and goat α-chicken-Alexa Fluor 488 (1:500, cat.no. A11039, Invitrogen) to better visualize the *sebox::EGFP* domain. After imaging, the explants were genotyped as described (Pomreinke *et al*, 2017) to identify homozygous mutants.

For immunostaining of pSmad5 upon simultaneous overactivation of BMP signaling and reduction in Nodal signaling activity, wildtype embryos were injected with 1 pg *bmp2b* and 80 pg *membrane-GFP* mRNA at the 1-cell stage and treated with 0.1% DMSO and 1 µM or 10 µM of the Nodal inhibitor SB-505124 (Byfield *et al*, 2004; Rogers *et al*, 2017), respectively, from 4-16-cell stage until 75% epiboly. Fixation and further processing for immunostaining was performed as described above.

### Embryo axis length measurements

To analyze embryo axis length upon BMP overactivation, Tg(*sebox::EGFP*) embryos were injected with 80 pg *membraneRFP* (for controls) or 30 pg *caAlk8* with 80 pg *membraneRFP* and fixed at bud stage in 4% PFA at 4°C overnight. To analyze embryo axis length upon BMP overactivation and simultaneous lowering of Nodal signaling levels, Tg(*sebox::EGFP*) embryos were injected with *membrane-RFP* or *H2A-mChFP* (for controls) or 1-5 pg *bmp2b* with *membrane-RFP* or *H2A-mChFP* and treated with 0.1% DMSO (control) or 1 µM SB-505124 (Byfield *et al*, 2004; Rogers *et al*, 2017) from 4-16 cell stage until bud stage and fixed at bud stage in 4% PFA at 4°C overnight. The embryos were washed in PBS + 0.1% Tween and mounted in 0.7% low melting point agarose in lateral view on glass bottom dishes (µ-Slide 4 Well, Ibidi) for imaging on a Zeiss LSM880 inverted microscope with a Zeiss Plan-Apochromat 10x objective.

### Quantitative real-time PCR (qRT-PCR)

Total RNA extraction of 10 explants prepared from wildtype, MZ*oep* or MZ*lefty1,2* embryos at 75% epiboly was performed using 0.5ml Trizol (Invitrogen) according to the manufacturer’s protocol. Genomic DNA removal was performed using the DNA-free DNA Removal Kit (Thermo Fisher Scientific) according to the manufacturer’s protocol. cDNA and negative control NO-RT cDNA reactions were performed using the iScript™ Reverse Transcription Supermix for RT-qPCR according to the manufacturer’s protocol with 500ng total RNA as starting material. Primers for qRT-PCR were checked for linear amplification by a concentration series of cDNAs. For the experiment, a 1:10 cDNA dilution was used. As a housekeeping gene for normalization, we used *elongation factor 1 α* (*EF1α*) (Miesfeld *et al*, 2015). Intron-spanning primers for *chrd* were designed using Primer3: 5’-CGA CTC TTC CAC CAA TCA CA-3’, 5’-CAG ATA CGC CGT ACC TTC AT-3’. The QPCR SYBR Green Mix (Thermo Scientific) was used for all qRT-PCRs, which were run on a Bio-Rad C1000 Thermal Cycler. Reactions were performed in triplicates.

### Image analysis

Image analysis was performed using Bitplane Imaris or Fiji (Schindelin *et al*, 2012).

### Analysis of explant morphology

For quantifying explant and mesendoderm morphogenesis in high-resolution live imaging timelapse movies, the onset of extension was defined as the timepoint after which a clear elongation of the explant could be seen in more than three subsequent frames on a 3D reconstruction in Bitplane Imaris. To quantify the shape of the explants, images were exported to Fiji and the explant circumference was outlined based on brightfield images to generate a binarized image of the explant and the background by thresholding. A reference axis was defined by drawing a 40 pixels wide box in the center of the explant. The morpholibJ geodisc distance map function was used in Fiji to calculate intensity values based on the distance of each pixel in the explant from the reference axis. These intensities were used to reconstruct the explant shape by measuring the intensity values of the furthest points from the axis on one side of the explant.

For length measurements of explants at specific developmental stages, the distance from the tip of the explant to the back was measured in Fiji using the Segmented Line Tool on bright-field side view images. Quantification of blastoderm explant morphology on bright-field side view images was performed as described previously (Schauer *et al*, 2020). In short, explants were considered extended if a clear indentation could be seen between the round and extended part of the explant. Explants were manually outlined to quantify explant circularity using the Circularity plugin in Fiji. Extended and not extended explants were pooled in this analysis. The length of the explant extension was quantified by measuring the distance from the tip of the explant to the indentation point with the Segmented Line tool from Fiji and normalized to the full explant length. Only explants where the extension was clearly visible were included in this analysis.

For measuring explant length upon local *caAlk8* overexpression, clones were categorized based on clone localization at bud stage into *mostly mesendodermal* (if less than 25% of the clone were outside of the *sebox*::*EGFP*-positive domain), *mostly ectodermal* (if less than 25% of the clone were inside of the *sebox*::*EGFP*-positive domain) or *ectodermal plus mesendodermal* clones (if at least 25% of the clone can be found in both domains). The segmented line tool was then used in Fiji to measure the length from the tip of the extension to the base of the extension and the length of the whole explant on maximum intensity projections based on brightfield images in side-view.

### Analysis of *sebox*::*EGFP* domain morphology

To approximate the proportion of mesendoderm within the extension of explants, the onset of explant extension was identified as described above. In Fiji, a SUM projection was made at the indicated timepoints after the onset of extension. The image was then binarized by thresholding and a median filter of 2 pixels width was applied to reduce noise. The circumference of the extension and base of the extension were outlined on the brightfield images using the freehand selection tool in Fiji. This mask was then applied to quantify the area fraction on the binarized *sebox::EGFP* image to measure the proportion of mesendodermal tissues in the extension. Note that for this quantification, explants where the ectoderm overlayed mesendoderm in the plane of the projection had to be excluded. To analyze the localization of the *sebox::EGFP* domain along the extension, images were binarized as described. A 50 pixel wide line was drawn in the middle of the explant from the tip of the extension towards the end of the explant, and the plot profile function was used to get intensity values along this axis.

The quantify the length and width of the *sebox::EGFP* domain over time, the Segmented Line Tool was used in Fiji to measure the extent of the GFP signal in the direction of the extension (length) and the extent of the GFP signal in the middle of the domain perpendicular to the extension (width). To quantify the length/width ratio of the *sebox::EGFP* domain upon BMP overactivation, length and width were similarly determined. The area of the *sebox::EGFP* domain was measured by manually outlining the GFP positive domain using the freehand selection tool in Fiji on Maximum intensity projections. The boundaries of the domain were confirmed by looking through individual sections to reduce information loss due to the projection.

### Analysis of clone dispersal

To automatically track clone cells in blastoderm explants in 3D based on their nuclear signal at a time resolution of less than 3 minutes, the Spot Detection Plugin from Imaris was employed. Tracking parameters were set using autoregressive motion and allowing a gap of 1 frame in the tracks and a maximum distance of 10 µm between subsequent timepoints. All tracks were manually verified, and incomplete tracks were either removed or manually completed to encompass the whole analysis timeframe. The tracks were divided into mesendodermal and ectodermal cells based on their location within the *sebox::EGFP* positive or negative domain, respectively, and their 3D coordinates were extracted for further analysis. We used a custom MATLAB script (Supplementary File 1) to calculate the mean distance of each cell to all clone cells in the direction of the extension and perpendicular to it in a 2D coordinate system. As explants were not stably positioned during imaging, the main explant axis was defined manually by placing a spot at the bottom of the analysis volume at the extension tip every 10 time frames and a spot which was placed at the round part of the explant opposite the extension. All spots were projected along their Z-axis, and the mean distance of each cell to other cells in the clone was measured either in the direction of the extension or perpendicular to it, relative to the initial time point. Notably, tracks that began at later time points were excluded from the analysis.

### Analysis of cell alignment

Cell alignment was measured as described previously (Williams & Solnica-Krezel, 2020b). In short, explants and embryos were oriented with the direction of explant elongation and anterior pole of the embryo pointing up, as identified based on short timelapse movies. Rotation of embryos was performed in 3D in Imaris in cases where the dorsal side of the embryo was not parallel to the imaging plane. The outlines of the cell body were drawn based on membrane or Lifeact (F-actin) signal on individual planes using the freehand selection tool in Fiji, and the angle of the longest axis of the best fit ellipse to the main explant or mediolateral embryonic axis was measured.

### Analysis of gene expression domains

The area of gene expression domains was determined by outlining the signal based on *in situ* hybridizations in Fiji using the freehand selection tool and normalized to the area of the whole explant. The length of the explants was measured using the freehand selection tool. To exclude the possibility that explants were lacking Nodal signaling altogether, only explants forming an elongation were considered for this analysis.

### Analysis of pSmad5 and pSmad2 nuclear localization

Signaling pathway activity profiles were determined as described previously using a custom MATLAB script (Schauer *et al*, 2020). In short, the Spot Detection function in Imaris was used to automatically determine nuclear coordinates based on DAPI signal. Nuclei were analyzed to a depth of 180 µm from the top for explants in side-view and at 60-120 µm from the animal pole for embryos. Enveloping layer nuclei, yolk syncytial layer nuclei and nuclei with inhomogeneous DAPI signal as well as clearly dividing cells were manually excluded before further analysis steps as described previously (Schauer *et al*, 2020). Reference points to calculate signaling activity profiles were determined by manually drawing a line of nuclei using the same Spot Detection tool on the side of highest pSmad5 intensity in explants and embryos. 3D coordinates for the reference points and all nuclei as well as mean fluorescence intensities for the nuclei were then extracted, and a previously published custom MATLAB script (Schauer *et al*, 2020) was used to calculate the geometric distance of all nuclei in the explant or embryo from the reference points upon projection along their Z axis.

Background subtraction was performed by manually placing 12 spots in the cytoplasm of cells located on the low pSmad5 side at the bottom of the analysis volume. For double stainings of pSmad5 and pSmad2, 12 spots were placed in the cytoplasm of cells located on either side at the bottom of the analysis volume. pSmad5 and pSmad2 intensities were normalized to the DAPI signal to correct for artifacts due to imaging depth and high intensity nuclei were manually re-checked to exclude potential artificially high nuclei due to very weak or inhomogeneous DAPI or dividing nuclei following the criteria for the initially described correction. To account for differences in explant and embryo size, the calculated distances were normalized to the maximum distance in each explant/embryo. To plot the normalized pSmad5 and pSmad2 profiles from the low pSmad5 signaling side, the data were binned into steps of 0.04 for explants and 0.02 for embryos, as this bin size corresponds to approximately 2 cell diameters, respectively, and averaged across the experimental replicates. The first and last 0.04 bin of explants and embryos were excluded since they contain only few nuclei due to the sample curvature. Note that for intensity measurements the same number of control and treated embryos or blastoderm explants have been analyzed per replicate in each graph to reduce the inherent variability due to processing and imaging on different experimental days between replicates. The intensities are shown relative to the average intensity in the first bin closest to the high pSmad5 and the high pSmad2 side of control samples.

To quantify the extent of the pSmad5 and *sebox::EGFP* positive domains along the explant axis, we performed maximum intensity projections from the middle plane of the explant to the top of the explant in Fiji. We then measured the distance from the back of the explant to the end of the pSmad5 positive domain, as assessed by the presence of clearly visible nuclear pSmad5 staining, in the center of the explant using the Line Tool. Similarly we determined the distance from the back of the explant to the start of the mesendoderm as assessed by the presence of *sebox::EGFP* signal and normalized to the overall length of the explant.

Categorizing pSmad5 profiles as graded or radial in control conditions or upon overexpression of 1pg *bmp2b* plus treatment with 1 µM or 10 µM SB-505124 was done by defining a radial distribution when nuclear pSmad5 levels appear uniformly elevated, while in graded conditions, a clear difference in pSmad5 levels along the prospective DV axis can be found. To quantify the ratio of DV pSmad5 levels, nuclei were identified based on the DAPI signal by the Spot Detection Tool in Imaris as described above. 10 nuclei were then randomly selected in the highest pSmad5 intensity domain (ventral) and lowest pSmad5 domain (dorsal) at 110-120 µm from the top of animally oriented embryos at 75% epiboly. In embryos where the low and high pSmad5 intensity domains were not clearly distinguishable due to the radial expansion of pSmad5 activity, nuclei were selected in randomly chosen opposing domains. The pSmad5 intensity was normalized to DAPI intensity and the ratio of dorsal to ventral nuclei was calculated.

### Analysis of embryonic axis length

The length of the embryonic axis was determined by measuring the distance from the front of the *sebox::EGFP* domain to the tailbud using the segmented line tool along the curvature of the embryo in Fiji on maximum intensity projections. Embryo height was determined by measuring the distance from the head domain of the embryo to the tailbud.

### Statistics

GraphPad Prism was used to perform statistical analysis and generate plots. The number of analyzed blastoderm explants or embryos (n) and experimental replicates (N) are stated in the figure legends. Error bars in box plots correspond to the minimum and maximum in the dataset. Error bars in graphs either correspond to ± S.D. or ± S.E.M. as indicated on the respective axis labels. All data were tested for normality before choosing the statistical test to assess significance using the D’Agostino–Pearson normality test. In Fig. S3I, the sample size was insufficient for the D’Agostino–Pearson normality test in one category and we thus used the Shapiro-Wilk test to assess normality. For comparison of two groups, we used a two-sided Student’s t test or a Mann-Whitney test, depending on the datasets showing normal distribution or not. For comparison of more than two sample groups, we used an ANOVA or Kruskal-Wallis test, depending on the datasets showing normal distribution or not. A correction for multiple comparisons was performed in such cases. The specific statistical tests as well as exact p-values are stated in the figure legends.

