## Supplementary File 1 for "Robust axis elongation by Nodal-dependent restriction of BMP signaling": CellDispersal.m

```
%changed output to absolute distance on parallel axis (from norm.)
%added center of mass

clearvars; close all;
[currentpath,~,~]=fileparts(mfilename('fullpath'));
addpath(currentpath);
%dbstop if error;

axisString='axis';
expSubString='clone';
ProjectinZdirection=true;
ProjectinZdirectionEvenForCloneScatterPlots=false;
showStartEndClone=true;
saveExcelOutput=true;
resultString='_Dist.xlsx';
resultCenterOfMassString=['_CenterOfMass'  resultString]; 

selectfiles=true;
if selectfiles
    workDir=uigetdir('J:\heisegrp\Alex\for robert','Select Data Folder');    
else
    workDir='B:\fs3-bifexchgrp\BIF_StaffSci\Robert\Alex\cellDispersal\data\20210318_wt_tracking_256cE_50_900Up - clone analysis';
end

cd(workDir);
allFiles=dir([expSubString '*.xls']);
cd(currentpath);

axisFile=fullfile(workDir, [axisString, '.xls']);
A=readtable(axisFile,'Sheet','Position','NumHeaderLines',1);
nTimepoints=height(A)-1;
apAxis1=zeros(nTimepoints,3);
apAxis2=zeros(nTimepoints,3);
t=zeros(nTimepoints,1);
for i=1:nTimepoints
    t(i)=A.Time(i+1);
    apAxis1(i,:)=[A.PositionX(1),A.PositionY(1),A.PositionZ(1).*(~ProjectinZdirectionEvenForCloneScatterPlots)];
    apAxis2(i,:)=[A.PositionX(i+1),A.PositionY(i+1),A.PositionZ(i+1).*(~ProjectinZdirectionEvenForCloneScatterPlots)];
end

X={};X2={};
minMaxTime=[];
nFiles=0;
for i=1:length(allFiles)
    if contains(allFiles(i).name,expSubString)
        nFiles=nFiles+1;
        titleString{nFiles}=allFiles(i).name;
        A=readtable(fullfile(workDir,allFiles(i).name),'Sheet','Position','NumHeaderLines',1);
        X{nFiles}=[A.PositionX,A.PositionY,A.PositionZ.*(~ProjectinZdirectionEvenForCloneScatterPlots),A.Time, A.TrackID-1E9];
        X2{nFiles}=[A.PositionX,A.PositionY,A.PositionZ,A.Time]; % saved for writing results. 
        minMaxTime=vertcat(minMaxTime,[min(A.Time) max(A.Time)]);
        if showStartEndClone
            figure
            title(titleString{nFiles});
            hold('on');
            scatter3(apAxis1(end,1),apAxis1(end,2),apAxis1(end,3),'O','SizeData',80);
            text(apAxis1(end,1),apAxis1(end,2),apAxis1(end,3),'a');
            scatter3(apAxis2(end,1),apAxis2(end,2),apAxis2(end,3),'O','SizeData',80);
            text(apAxis2(end,1),apAxis2(end,2),apAxis2(end,3),'b');
            x=X{nFiles};
            x1=x(x(:,4)==minMaxTime(nFiles,1),:);
            x2=x(x(:,4)==minMaxTime(nFiles,2),:);
            scatter3(x1(:,1),x1(:,2),x1(:,3),'x','MarkerEdgeColor','green');
            scatter3(x2(:,1),x2(:,2),x2(:,3),'x','MarkerEdgeColor','blue');

            hold('off');
        end
    end
end

if ProjectinZdirection
    apAxis1(:,3)=0;
    apAxis2(:,3)=0;
    for i=1:nFiles
        X{i}(:,3)=0;
    end
end

% debug
% figure
% hold('on');
% scatter3(apAxis1(1,1),apAxis1(1,2),apAxis1(1,3));
% plot3(apAxis2(:,1),apAxis2(:,2),apAxis2(:,3),'LineWidth',3);
% plot3(smooth(apAxis2(:,1)),smooth(apAxis2(:,2)),smooth(apAxis2(:,3)),'LineWidth',3);
%
% [d, p, pNorm]=distanceFromProjectionontoLine(X{1}(1,1:3),apAxis1(1,:),apAxis2(1,:));
%
% figure;
% hold('on');
% scatter3(X{1}(1,1),X{1}(1,2),X{1}(1,3));
% scatter3(apAxis1(1,1),apAxis1(1,2),apAxis1(1,3),'x');
% scatter3(apAxis2(1,1),apAxis2(1,2),apAxis2(1,3),'O');
% plot3([apAxis1(1,1),apAxis2(1,1)],[apAxis1(1,2),apAxis2(1,2)],[apAxis1(1,3),apAxis2(1,3)]);
%
% y=(apAxis2(1,:)-apAxis1(1,:)).*pNorm+apAxis1(1,:);
% plot3([y(1),X{1}(1,1)],[y(2),X{1}(1,2)],[y(3),X{1}(1,3)]);
% plot3([y(1),apAxis1(1,1)],[y(2),apAxis1(1,2)],[y(3),apAxis1(1,3)],'LineWidth',2);


%distances of the cells with respect to axis
verticalAndParallelDistances=cell(nFiles,1);
for n=1:nFiles
    k=length(X{n});
    dVertical=nan(k,1);
    dParallel=nan(k,1);
    dParallelNorm=nan(k,1);
    for i=1:k
        [~,minIndex]=min(abs(t-X{n}(i,4))); % closest time point
        [dVertical(i), dParallel(i) ,dParallelNorm(i)]= distanceFromProjectionontoLine(X{n}(i,1:3),apAxis1(minIndex,:),apAxis2(minIndex,:));
    end
    figure
    subplot(1,2,1);
    histogram(dParallel,20);
    title([titleString{n}  ': parallel'],'Interpreter','none' );
    subplot(1,2,2);
    histogram(dVertical,20);
    title([titleString{n}  ': vertical'],'Interpreter','none' );
    figure;
    subplot(1,2,1);
    scatter(X{n}(:,4),dParallel, 'x');
    xlabel('time');
    title([titleString{n}  ': parallel'],'Interpreter','none' );
    subplot(1,2,2);
    scatter(X{n}(:,4),dVertical,'x');
    xlabel('time');
    title([titleString{n}  ': vertical'],'Interpreter','none' );
    verticalAndParallelDistances{n}=[dParallel,dVertical];

end


% distances of the cells with respect relative to each other
% duplicate calculation so it's easier to change the axis etc. 
centerOfMass=cell(nFiles,1);
for n=1:nFiles

    dParallel=verticalAndParallelDistances{n}(:,1);
    dVertical=verticalAndParallelDistances{n}(:,2);

    k=length(X{n});
    dVerticalOrigin=nan(k,1);
    dParallelOrigin=nan(k,1);
    dVerticalmeanDistToOtherCells=nan(k,1);
    dParallelmeanDistToOtherCells=nan(k,1);
    dVerticalmaxDistToOtherCells=nan(k,1);
    dParallelmaxDistToOtherCells=nan(k,1);
    dParallelDistToCenterofMass=nan(k,1);
    dVerticalDistToCenterofMass=nan(k,1);
    dVerticalmeanDistToOtherCellsNormT1=nan(k,1);
    dParallelmeanDistToOtherCellsNormT1=nan(k,1);

    tracks=unique(X{n}(:,5));
    timePoints=unique(X{n}(:,4));

    % center of mass calculation
    dVerticalCenterOfMass=nan(numel(timePoints),1);
    dParallelCenterOfMass=nan(numel(timePoints),1);
    for i=1:numel(timePoints)
        idx=find(X{n}(:,4)==timePoints(i));
        dParallelCenterOfMass(i,1)=mean(dParallel(idx));
        dVerticalCenterOfMass(i,1)=mean(dVertical(idx));
    end
    centerOfMass{n}=[dParallelCenterOfMass,dVerticalCenterOfMass,timePoints];

    % debug
    %     figure;
    %     hold('on');
    %     for i=1:numel(tracks)
    %         tx=tracks(i);
    %         plot(X{n}(X{n}(:,5)==tx,1),X{n}(X{n}(:,5)==tx,2));
    %     end
    %     hold('off');

    x0=nan(numel(tracks),3); % starting point of the track
    dPx0=nan(numel(tracks),1); % starting distance rel. Axis (paralle)
    dVx0=nan(numel(tracks),1); % starting distance rel. Axis (vertical)
    for i=1:numel(tracks)
        tx=tracks(i);
        x1Track=X{n}(X{n}(:,5)==tx,:);
        x0(i,:)=x1Track(x1Track(:,4)==min(x1Track(:,4)),1:3); % starting point of the track
        [~,minIndex]=min(abs(t-X{n}(i,4))); % closest time point
        [dVx0(i), dPx0(i), ~]= distanceFromProjectionontoLine(x0(i,:),apAxis1(minIndex,:),apAxis2(minIndex,:));
    end
    for i=1:k
     
        dVerticalOrigin(i)=dVertical(i)-dVx0(tracks==X{n}(i,5));
        dParallelOrigin(i)=dParallel(i)-dPx0(tracks==X{n}(i,5));
        
        % mean distance
        tC=X{n}(i,4);  % current time point
        sameT=find(X{n}(:,4)==tC);
        sameT(sameT==i)=[];
        if ~isempty(sameT)
           dVerticalmeanDistToOtherCells(i)=mean(abs(dVertical(sameT)-dVertical(i))); 
           dParallelmeanDistToOtherCells(i)=mean(abs(dParallel(sameT)-dParallel(i)));
           dVerticalmaxDistToOtherCells(i)=max(abs(dVertical(sameT)-dVertical(i))); 
           dParallelmaxDistToOtherCells(i)=max(abs(dParallel(sameT)-dParallel(i)));
        end

        % dist to center of mass
        dParallelDistToCenterofMass(i)=abs(dParallel(i)-dParallelCenterOfMass(timePoints==tC));
        dVerticalDistToCenterofMass(i)=abs(dVertical(i)-dVerticalCenterOfMass(timePoints==tC));
    
       % normalized distance to  t=1
       currentTrackId=X{n}(i,5);
       idT1=find(X{n}(:,5)==currentTrackId & X{n}(:,4)==t(1)); % first time point
       if isempty(idT1)
          
          %discard
          dVerticalmeanDistToOtherCellsNormT1(i)= nan;
          dParallelmeanDistToOtherCellsNormT1(i)= nan;

       else
          dVerticalmeanDistToOtherCellsNormT1(i)= dVerticalmeanDistToOtherCells(i)/ dVerticalmeanDistToOtherCells(idT1);
          dParallelmeanDistToOtherCellsNormT1(i)= dParallelmeanDistToOtherCells(i)/ dParallelmeanDistToOtherCells(idT1);
       end

      

    end

    figure
    subplot(1,2,1);
    histogram(dParallelOrigin,20);
    title([titleString{n}  ': parallel distance to start point'],'Interpreter','none' );
    subplot(1,2,2);
    histogram(dVerticalOrigin,20);
    title([titleString{n}  ': vertical distance to start point'],'Interpreter','none' );
    figure;
    subplot(1,2,1);
    scatter(X{n}(:,4),dParallelmeanDistToOtherCells, 'x');
    xlabel('time');
    title([titleString{n}  ': mean parallel distance to other cells'],'Interpreter','none' );
    subplot(1,2,2);
    scatter(X{n}(:,4),dVerticalmeanDistToOtherCells,'x');
    xlabel('time');
    title([titleString{n}  ': mean vertical distance to other cells'],'Interpreter','none' );

    figure;
    subplot(1,2,1);
    scatter(X{n}(:,4),dParallelmaxDistToOtherCells, 'x');
    xlabel('time');
    title([titleString{n}  ': max parallel distance to other cells'],'Interpreter','none' );
    subplot(1,2,2);
    scatter(X{n}(:,4),dVerticalmaxDistToOtherCells,'x');
    xlabel('time');
    title([titleString{n}  ': max vertical distance to other cells'],'Interpreter','none' );

    figure;
    subplot(1,2,1);
    scatter(X{n}(:,4),dParallelmeanDistToOtherCellsNormT1, 'x');
    xlabel('time');
    title([titleString{n}  ': Norm T1 parallel distance to other cells'],'Interpreter','none' );
    subplot(1,2,2);
    scatter(X{n}(:,4),dVerticalmeanDistToOtherCellsNormT1,'x');
    xlabel('time');
    title([titleString{n}  ': Norm T1 vertical distance to other cells'],'Interpreter','none' );


    vertParaOriginAndMeanDistances{n}=[dParallelOrigin, dVerticalOrigin, abs(dParallelOrigin), abs(dVerticalOrigin),dParallelmeanDistToOtherCells, ...
                                       dVerticalmeanDistToOtherCells, dParallelmaxDistToOtherCells, dVerticalmaxDistToOtherCells, ...
                                       dParallelDistToCenterofMass, dVerticalDistToCenterofMass, dParallelmeanDistToOtherCellsNormT1, dVerticalmeanDistToOtherCellsNormT1];

end


if saveExcelOutput
    for i=1:nFiles
        outname=fullfile(workDir,[titleString{i}(1:end-4) resultString]);
        A=[X2{i} verticalAndParallelDistances{i} vertParaOriginAndMeanDistances{i}];
        disp(['writing ' outname]);
        writecell({'X','Y','Z','t','parallel distance','vertical distance','parallel distance from origin','vertical distance from origin',...
                   'abs. parallel distance from origin','abs. vertical distance from origin',...  
                   'mean parallel distance to other cells','mean vertical distance to other cells',...
                   'max parallel distance to other cells','max vertical distance to other cells',...
                   'parallel distance to center of mass', 'vertical distance to center of mass', ...
                   'mean parallel distance to other cells normalized to t=1','mean vertical distance to other cells normalized to t=1'},outname);
        writematrix(A,outname,'Sheet',1,'Range','A2');

        outname=fullfile(workDir,[titleString{i}(1:end-4) resultCenterOfMassString]);
        A=centerOfMass{i};
        writecell({'center of mass: parallel distance','center of mass: vertical distance','t', },outname);
        writematrix(A,outname,'Sheet',1,'Range','A2');

    end
end
```
