## Supplementary File 1 for "Robust axis elongation by Nodal-dependent restriction of BMP signaling": distanceFromProjectionontoLine.m

```
function [d, p, pNorm] = distanceFromProjectionontoLine(pt, v1, v2)
      
a = v1 - v2;
b = pt - v2;
d = norm((cross(a,b)) / norm(a)); % distance from line
u=v2-v1;
v=pt-v1;
p=dot(v,u)/norm(u);
pNorm=p/norm(a);

end
```
